# Disrupting aberrant EGFR catalytic trimers reverses T790M gefitinib resistance

**DOI:** 10.64898/2026.09.24.754095

**Authors:** Selene K. Roberts, Ioannis Galdadas, Nicola Piasentin, Sarah R. Needham, Benjamin M. Davis, Laura C. Zanetti-Domingues, Rico C. H. Man, David T. Clarke, Andrew H. A. Clayton, Daniel J. Rolfe, Gilbert O. Fruhwirth, Francesco L. Gervasio, Marisa L. Martin-Fernandez

**Affiliations:** Central Laser Facility, UKRI-STFC Rutherford Appleton Laboratory, Didcot, Oxfordshire, UK; School of Pharmaceutical Sciences, University of Geneva, Geneva, CH; ISPSO, University of Geneva, Geneva, CH; Atomistic Simulations, Italian Institute of Technology, Genova, IT; Imaging Therapies and Cancer Group, Comprehensive Cancer Centre, School of Cancer and Pharmaceutical Sciences, King’s College London, London, UK; Optical Sciences Centre, School of Science, Computing and Emerging Technologies, Swinburne University of Technology, Melbourne, Australia

## Abstract

Epidermal growth factor receptor (EGFR) mutations drive up to 50% of non-small-cell lung cancers (NSCLC). Although tyrosine kinase inhibitors (TKIs) like gefitinib initially offer substantial clinical benefit, the emergence of acquired, on-target resistance inevitably limits long-term remission. Beyond structural changes that alter ATP-pocket affinity, EGFR oligomerization drives this resistance, though the underlying mechanisms remain unclear. Here we show that progressive, TKI-resistant secondary and tertiary EGFR-mutant variants assemble on the cell surface into higher-order, ligand-free oligomers organized around pathological catalytic trimers. In contrast, gefitinib-sensitive variants assemble into oligomers where the catalytic units remain as canonical dimers, like those found in wild-type oligomers. Genetically disrupting these pathological trimers into dimers via a single-point mutation rewires downstream signaling, decelerates tumor progression, and reverses gefitinib resistance *in vivo*. Conversely, genetic engineering of dimers into trimers reinstates aggressive tumor growth. These findings expose a structural vulnerability unique to acquired mutations, demonstrating that selectively targeting these pathological intra-oligomer trimers can overcome resistance, and provide a blueprint for protein-protein interface modulation strategies that spare wild-type signaling.

## INTRODUCTION

Activating mutations within exon 18–21 of the *EGFR* gene are oncogenic drivers in non-small-cell lung cancer (NSCLC), a disease with a profound global burden^1^. For decades, successive generations of anti-EGFR tyrosine kinase inhibitors (TKIs) attempted to outpace the inevitable emergence of on-target resistance. Despite these therapeutic advances, the median maximum overall survival remains approximately 3.3 years^2^. This poor prognosis is particularly concerning given the global rising incidence of NSCLC among young, non-smoking adults, a demographic where *EGFR* mutations represent the predominant oncogenic driver^3^.

First-generation tyrosine kinase inhibitors (TKIs), such as gefitinib, initially transformed the clinical outcomes of patients presenting with classical *EGFR* mutations^4,5^. These variants constitute ~85-90% of *EGFR*-mutated NSCLC cases, primarily comprising exon 19 deletions (e.g. delE746_A750, or ΔELREA) and the exon 21 L858R single point mutation. However, after approximately 10 months, patients develop resistance driven by secondary *EGFR* mutations, most notably the exon 20 T790M ‘gatekeeper’ substitution^6,7^. Subsequent drug iterations, including the third-generation osimertinib, which targets the T790M mutation^8^, continue to drive the Darwinian emergence of tertiary *EGFR* mutations. This evolutionary pressure suggests that sequential small-molecule drug designs will inevitably face recurring, on-target resistance.

To disrupt this predictable cycle of resistance, structural insights are increasingly important for therapeutic discovery; however, structural data have historically been leveraged for post hoc rationalization^9^. More recently, biophysical studies have linked higher-order receptor oligomerization to resistant phenotypes^10,11^, yet the precise structural mechanisms remain obscure. Here, by integrating molecular dynamics (MD) simulations and super-resolution imaging we identified the structure alterations within higher-order oligomers that drive resistance to TKIs. In combination with targeted proteomics, biochemical and biophysical assays, and optical and electron microscopies, we reveal how these structural alterations rewire downstream signaling networks and how reversing them resensitized T790M-mutant tumors to gefitinib. Ultimately, our framework establishes an orthogonal paradigm for rational drug design and clinical management in resistant NSCLC.

### Simulations predict oncogenic and autoinhibited EGFR’s kinase domain trimers

In non-cancerous cells, ligand binding tightly controls formation of the canonical asymmetric kinase dimer (Asym_k_-dimer) of wild-type EGFR (EGFR^WT^)^12^. Asymmetric dimerization drives a 500-fold increase in ATP-catalysis relative to monomeric EGFR, enabling autophosphorylation and downstream signaling^13^. Conversely, oncogenic EGFR-mutants, like EGFR^L858R^, constitutively stabilize the Asym_k_-dimer to promote tumorigenesis^14^. Interestingly, the secondary resistant variant EGFR^T790M^ disfavors Asym_k_-dimer formation due to increased disorder within the *α*C-helix^15^.

Emerging evidence links ligand-independent, higher-order oligomerization to TKI resistance^10,16^. Assembly of these oligomers is governed by the evolutionarily important, tethered conformation of the EGFR ectodomain^17^. We previously proposed that the weak Asym_k_-dimer of EGFR^T790M^ assembles within such ligand-independent oligomers^11^ (Fig. 1) and hypothesized that a backbone-to-backbone interface (BbBbI) with an adjacent monomer strengthened this dimer. Because similarly weakened Asym_k_-dimers occur in other NSCLC EGFR-mutants^13,18^, we reasoned that this interface may act as a compensatory mechanism, stabilizing fragile Asym_k_-dimers across different variants.

**Figure 1.**
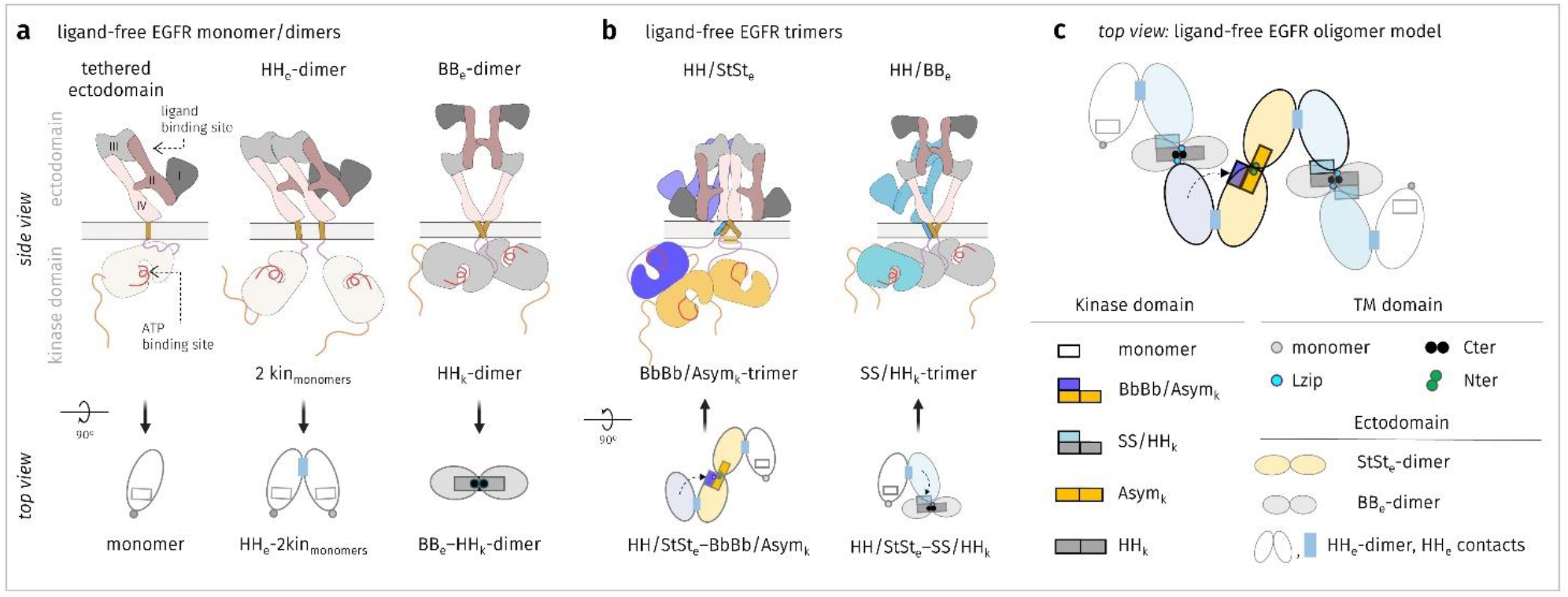
Schematic of catalytic and autoinhibited components of ligand-independent EGFR oligomers. **a**, Side and top views of the EGFR monomer and two autoinhibited dimers that EGFR can adopt at the plasma membrane: a head-to-head ectodomain dimer (HH_e_-dimer) that exhibits a tethered conformation^17^ and is linked to two uncoupled kinase monomers (2Kin-monomers)^15^, and a back-to-back ectodomain dimer linked to a head-to-head kinase dimer (BB_e_–HH_k_-dimer)^60^. This oligomer model was validated by mapping probe-binding sites using triangular geometric constraints from a two-dimensional version of FLImP^11^. **b**, Two HH_e_-dimer units combine to form a stalk-to-stalk ectodomain dimer (StSt_e_-dimer) coupled to an asymmetric kinase dimer (Asym_k_-dimer), the latter stabilized by forming a trimer via a backbone-to-backbone kinase interface (BbBbI) with an unengaged kinase monomer of a HH_k_-dimer. Unengaged kinase monomers can also interact with an adjacent BB_e_–HH_k_-dimer via a side-to-side interface (SSI) to form an autoinhibited trimer. Note that the ectodomains of individual HH_e_-dimers are colored according to the interaction that their kinase domains engaged in. One HH_e_ interface and the unengaged kinase monomer are not shown in the side views. **c**, Top view summarizing the ectodomain, transmembrane, and kinase interfaces present in the full ligand-free oligomer model.

Even transient stabilization of the Asym_k_-dimer by the BbBbI would require kinase domains to assemble trimers where a peripheral monomer is attached to the Asym_k_-dimer via a BbBbI (BbBb/Asym_k_-trimer). We used MD simulations to quantify these putative trimer stabilities. As a positive control, we used the exon 20 loop insertion variant D770_N771InsNPG (EGFR^InsNPG^), which exhibits weak asymmetric dimerization^18^ but possesses inserted residues within the core of the BbBbI that make this interface stable^11^ (Fig. 2a). We compared this control against two classical mutants (EGFR^ΔELREA^ and EGFR^L858R^ and three TKI-resistant variants. These include two secondary mutants, EGFR^T790M^ and the double mutant EGFR^L858R/T790M^ (EGFR^L/T^), which is commonly found in patients after receiving first-generation TKIs^7^, and the tertiary triple-mutant EGFR^L858R/T790M/C797S^ (EGFR^L/T/C^), which drives clinical resistance to third-generation osimertinib^19^.

**Figure 2.**
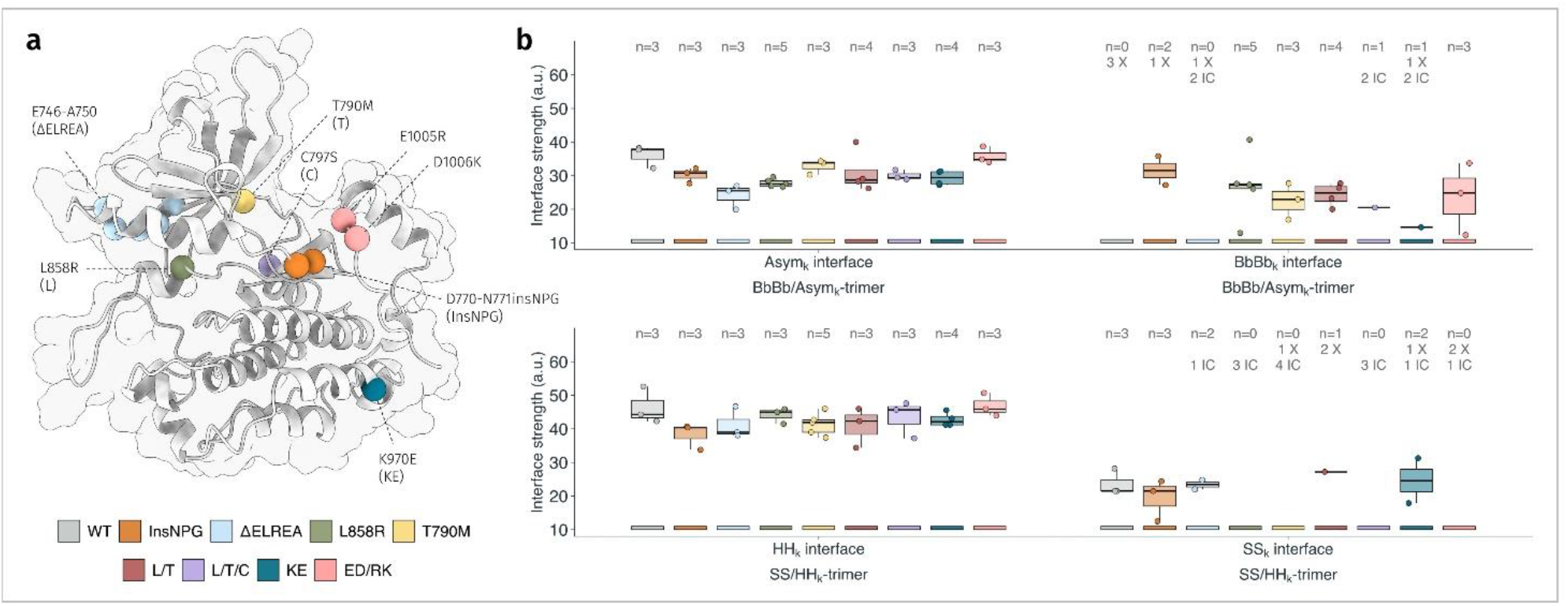
EGFR variants forms active and inactive stable kinase trimers. **a**, Kinase domain monomer with each mutation mapped as colored spheres, clustering around regions that are critical for kinase activation and dimerization. **b**, Interface strength (arbitrary units) measured across the four kinase interfaces present in the trimeric assemblies (Asym and BbBb interfaces of the BbBb/Asym_k_-trimer, top; HH and SS interfaces of SS/HH_k_-trimer, bottom) for each mutation. Mutations perturb these interfaces differently: some leave an interface largely unaffected, others breaking it entirely (X), and others substantially reshaping it while still holding the monomers together, in which case strength cannot be accurately quantified (IC); see Methods for details on the IC and X classifications. *n* indicates number of independent replicas used to quantify each interface (Details in Methods, Supplementary Fig. 1, and Supplementary notes 1-3).

In simulations of EGFR^WT^, starting from pre-assembled trimers, the monomer engaging via the BbBbI rapidly detached, leaving the Asym_k_-dimer behind (Fig. 2b, detachment reported as X). Conversely, the BbBbI monomer of EGFR^InsNPG^ remained bound, yielding the most stable BbBb/Asym_k_-trimers among all simulated conditions. Structurally stable BbBb/Asym_k_-trimers occurred across all oncogenic variants except EGFR^ΔELREA^, for which the BbBbI monomer remained physically attached but deviated substantially from the BbBbI geometry (interface change, IC). Because the Asym_k_-dimer core remained intact across all mutant simulations, our data suggest that the BbBbI monomer is dispensable to maintain the Asym_k_-dimer once formed. Instead, it acts as a structural scaffold that lowers the energetic barrier for initial asymmetric assembly. This scaffolding role may be enhanced by mutations that pre-organize monomeric structural elements, such as stabilizing the secondary structure of the *α*C-helix, which is known to favor Asym_k_-dimer assembly^12^.

Kinase monomers also establish a side-to-side interface (SSI) with a neighboring, autoinhibited head-to-head kinase dimer (HH_k_-dimer)^11,20^ (Fig. 1). This SS/HH_k_-trimer serves a dual role: it reinforces the leucine-rich transmembrane zipper interactions (Lzip) between oligomer constituents, while simultaneously depleting the local pool of available BbBbIs. Consequently, competition between BbBbIs and SSIs balances higher kinase activation from the BbBbIs with lower activation and a larger oligomer size from the SSIs^11^.

To determine whether NSCLC mutants exploit SSI destabilization to promote activation, we evaluated the stability of a putative inactive SS/HH_k_-trimer. While the SSI of EGFR^WT^ remained stable, this interface was destabilized in oncogenic mutants, even though the monomers engaging via the SSI remained attached in almost all replicates (Fig. 2b). This highlights a limitation of our computational approach in quantifying the interface strength of dimers that deviate substantially from their initial conformation. Nonetheless, our data below support the autoinhibitory nature of the SSI.

### Active trimers are associated with TKI resistance

Photobleaching imaging correlation spectroscopy^21^ revealed that EGFR^WT^ and its oncogenic variants assembled ligand-free cell surface oligomers (Extended Data Fig. 1). We analyzed these oligomers employing fluorescence localization imaging with photobleaching (FLImP), a ~2 nm resolution microscopy that uses a Bayesian framework to infer the most likely unique separations within an underlying structure^11,15,22^. FLImP analysis returned separations consistent with those predicted by the model in Fig. 1 (Extended Data Fig. 2). To investigate kinase trimer formation within these oligomers, we leveraged the principle of transmembrane structural coupling, whereby intracellular kinase interactions stabilize the local extracellular oligomer architecture^23^, which in turn reinforces specific separations between the ectodomains^11^ (Figs. 3a, 3b). This coupling enabled us to probe kinase domain interactions by non-disruptively labeling the EGFR ectodomains instead (red dots), a strategy that minimizes artefacts without disrupting the intracellular interactions of interest^11^. Using modeling and simulations, we estimated the theoretical uncertainties in the ectodomain separations coupled to the different kinase interactions (Fig. 3a, Extended Data Fig. 3).

**Figure 3.**
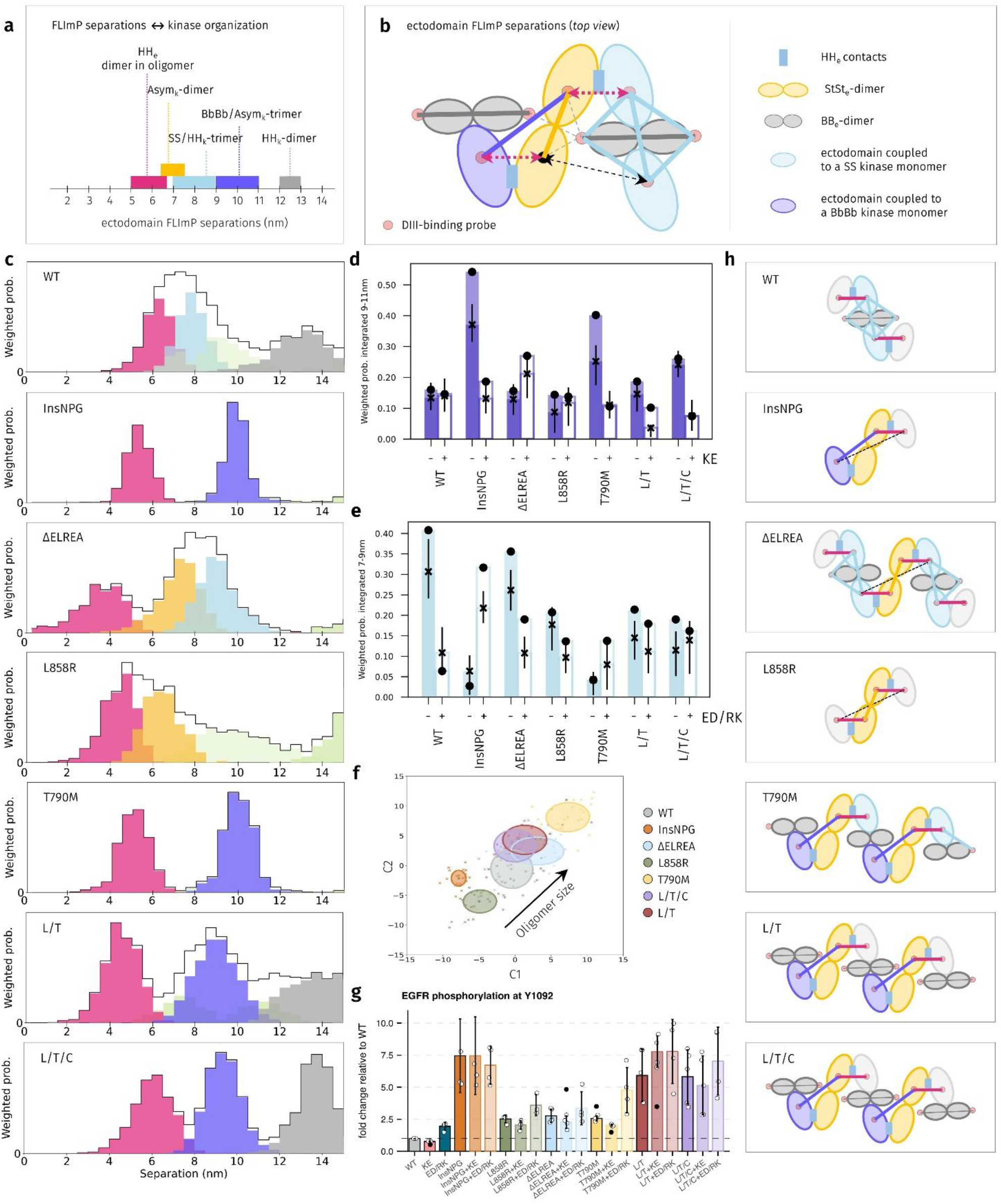
TKI-resistant and -permissive NSCLC mutations assemble distinct kinase trimer types. **a**, Separation ranges color-coded by kinase interaction. Boundaries derived from two structurally identical oligomer models, with probe separations in HH_e_-dimers set to 5 nm and 6.7 nm (Extended data Fig. 3). **b**, Top-view of an oligomer within the plane of the plasma membrane (kinase interactions not shown). Orange: Stalk-to-stalk ectodomain dimer linked to an asymmetric kinase dimer (StSt_e_–Asym_k_-dimer). Purple: protomer donating a BbBb kinase to form a BbBb/Asym_k_-trimer. Dark grey: back-to-back ectodomain dimers linked to head-to-head kinase dimers (BB_e_–HH_k_-dimers). Blue: ectodomains linked to kinases forming side-to-side interfaces with HH_k_-dimers (SS/HH_k_-trimers). Red circles: Affibody-CF640R probes labeling ectodomain DIII^11^. Black dot: Inaccessible probe-binding site when the BbBb/Asym_k_-trimer forms. Continuous colored lines: ectodomain separations stabilized by matching kinase interfaces (as in a). Dashed lines: pink, variable separations in HH_e_-dimers not stabilized by kinase interactions; black, separation commensurate with separations (~9-11 nm) stabilized by the BbBb/Asym_k_-trimer; gray, short separations (~4-6 nm) present when inner binding sites in BB_e_–HH_k_-dimers are accessible. **c**, FLImP analysis of the 100 highest-resolution empirical posterior separation distributions between Affibody-CF640R pairs for EGFR^WT^ and variants chosen from ~1,000-2,000 per variant. Peaks show abundance-weighted probability distributions of individual decomposed components, with colors reflecting association to a specific kinase interaction (components in pale green are not associated). Black lines show marginalized posterior distributions (the sum of the abundance-weighted peaks). Zoomed out 0-70 nm datasets are in Extended data Fig. 4. Supplementary Fig. 2. summarizes this analysis. **d, e** Quantification of separations in the 9–11 nm (with and without the KE mutation) and the 7-9 nm range (with and without the ED/RK mutations), showing FLImP abundance-weighted decomposition curves^11^ summed over the given ranges. Circles: values for original decompositions; crosses and error bars: mean and 20—80% confidence interval from decompositions on bootstrap-resampled FLImP measurements^11^. (FLImP decompositions used for these calculations are in Extended data Fig. 4). **f**, Relative oligomer sizes (black arrow marks the direction of size increase) calculated using Wasserstein multidimensional scaling (MDS) analysis of the FLImP decompositions in Extended data Fig. 4. The MDS metric maps similarities and differences across 21 separation sets of different conditions (one main plus 20 bootstrap-resampled decompositions). Plot axes are components C1 and C2 capturing the largest and second-largest orthogonal variance, respectively. Uncertainty ellipses represent 1SD. For more details see Supplementary Fig. 3. **g**, Western blot quantifications of ligand-independent phosphorylation in transfected CHO cells (n = 3 blots). Outliers identified using the Tukey 1.5×IQR rule (black circles) were excluded from bar heights and error bars (mean ±1SD). (Blots shown in Extended data Fig. 6a). **h**, Schematic top-view models depicting polymorphisms of the variants.

Within the 0–15 nm range relevant to local transmembrane structural coupling, FLImP decompositions show components that approximate the predicted positions, albeit with a broader distribution due to the method’s ~2 nm resolution (Figs. 3a, 3c). To determine whether the components spanning the 9-11 nm range reported stable BbBb/Asym_k_-trimers, as suggested in Fig. 3a (purple), we collected additional FLImP data to quantify separations in this range with and without the unnatural kinase domain mutation K970E (KE) (Extended Data, Fig. 4), which MD simulations predicted to disrupt the BbBb/Asym_k_-trimer (Fig. 2b). Consistent with simulations, the KE mutation reduced the highest component in the 9–11 nm range, observed for the BbBb/Asym_k_-trimer-positive control EGFR^InsNPG^, by approximately two-thirds (Figs. 3c, 3d). Similarly, introducing the KE mutation into EGFR^T790M^, EGFR^L/T^ and EGFR^L/T/C^ reduced their 9–11 nm peaks by one-half to two-thirds (Figs. 3c, 3d). Given the known oligomer architecture, these results demonstrate that EGFR^T790M^, EGFR^L/T^, and EGFR^L/T/C^ form stable BbBb/Asym_k_-trimers that are destabilized by the KE mutation, thereby validating the MD simulations for these variants.

**Figure 4.**
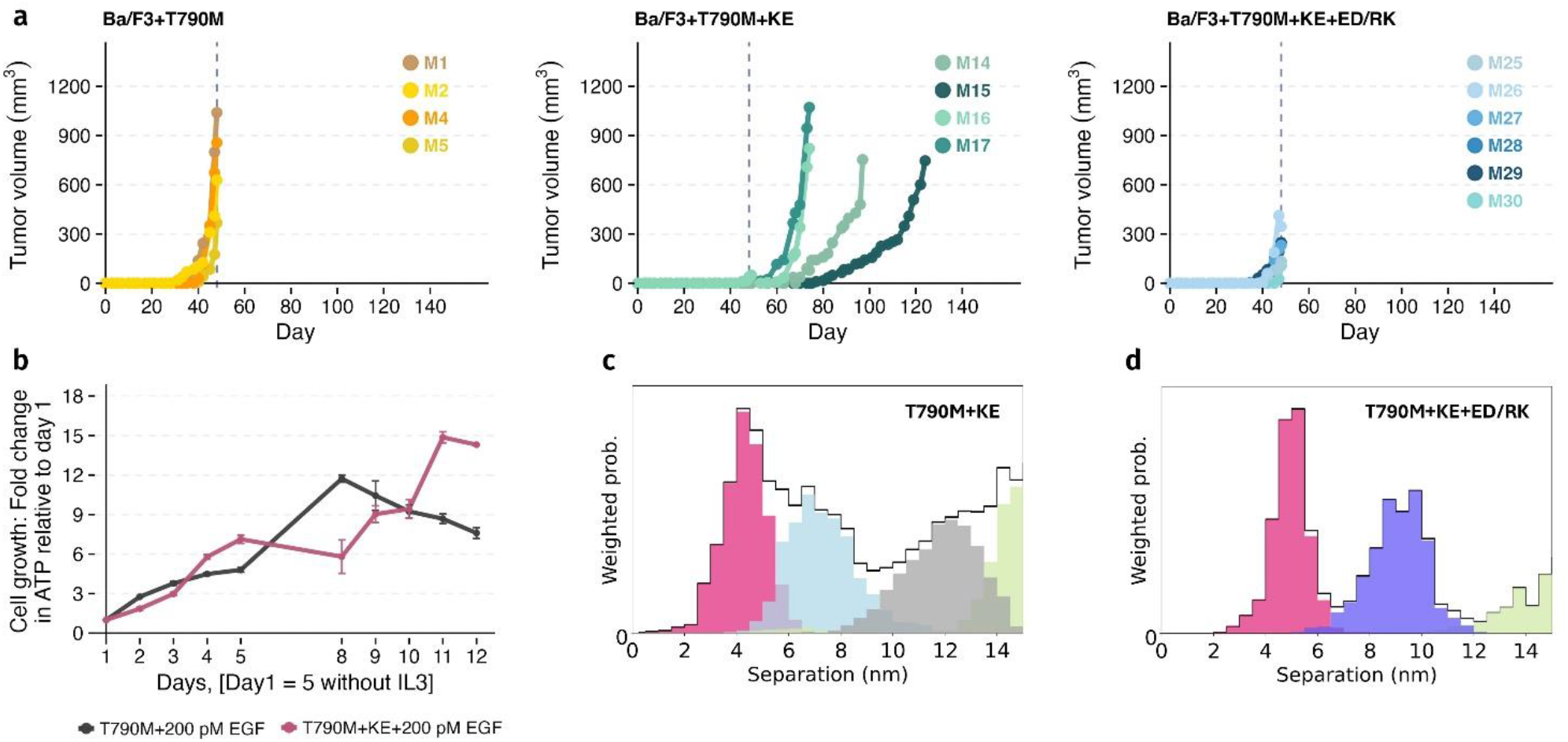
The transition from active trimers to dimers regulates tumor growth. **a**, *In vivo* tumor growth kinetics in NOD scid gamma (NSG) mice bearing subcutaneous xenografts of Ba/F3 cells expressing EGFR^T790M^, EGFR^T790M+KE^, or EGFR^T790M+KE+ED/RK^. Tumor volume was measured longitudinally using calipers. *n* = 6 biologically independent animals per cohort; four animals were censored (left and middle). Further details in Extended data Fig. 7. The vertical dashed line indicates day 48, when the first cohort reached the end point and was culled. Immunohistochemistry images of sections of these tumours are in Extended data Fig. 8a. **b**, *In vitro* growth curves of Ba/F3 cells stably expressing EGFR^T790M^ or EGFR^T790M+KE^. Cells were transfected using the PiggyBac system^11^ and fluorescence-activated cell sorted to minimize expression heterogeneity. Cells were deprived of IL-3 for five days (Days = –4 to 0), and 200 pM EGF was added at day 1. **c, d**, FLImP analysis of the highest resolution 100 empirical posterior separation distributions for EGFR^T790M+KE^ and EGFR^T790M+KE+ED/RK^ determined between Affibody-CF640R pairs. Peaks show abundance-weighted probability distributions of individual decomposed components, with colors reflecting association to a specific kinase interaction (components in pale green are not associated). Black lines show marginalized posterior distributions for each condition, the sum of the abundance-weighted peaks. A summary of this analysis is in Supplementary Fig. 2.

The KE mutation does not impact significantly the 9–11 nm separations for EGFR^WT^ and EGFR^ΔELREA^, consistent with predictions by MD simulations of BbBb/Asym_k_-trimer instability for EGFR^WT^ and EGFR^ΔELREA^ (Figs. 2b, 3c, 3d). The KE mutation has no effect on EGFR^L858R^ either, indicating that stable BbBb/Asym_k_-trimers are not abundant enough to be affected by this mutation. Contrary to this, MD simulations predicted stable BbBb/Asym_k_-trimers for EGFR^L858R^. This discrepancy likely arises because simulations initiate from a pre-assembled kinase trimer, while the absence of an effect from the KE mutation shows that EGFR^L858R^ BbBb/Asym_k_-trimers lack the free energy to significantly assemble spontaneously *in situ* in the first place. Consequently, we assigned the poorly resolved EGFR^L858R^ component centered at ~10 nm (pale green, Fig. 3c) to commensurate separations in the oligomers that are not coupled across the membrane with kinase interactions (black dashed arrow, Fig. 3b). These separations would be expected to return shallower, broader components, consistent with our observations. Likewise, for the KE-insensitive, split peak in EGFR^WT^ (pale green, Fig. 3c). These assignments are validated below.

### TKI-resistant mutations downregulate autoinhibited trimer conformations

FLImP decompositions also revealed components spanning the 7–9 nm ectodomain separations, where conformational coupling with the autoinhibited SS/HH_k_-trimers is expected (Figs. 3a, 3c light blue). To validate this assignment, we introduced the E1005R/D1006K (ED/RK) double mutation into the C-terminus of EGFR adjacent to the electrostatic hook that stabilizes the HH_k_-dimer^20^. Although we initially hypothesized that ED/RK would disrupt the HH_k_-dimer, thereby preventing SS/HH_k_-trimer formation, MD simulations instead predicted that ED/RK inhibits the SSI (Fig. 2a). Consistent with this prediction, ED/RK decreased the 7–9 nm separations for EGFR^WT^ and EGFR^ΔELREA^ (Fig. 3e), for which MD simulations predicted stable SS/HH_k_-trimers.

While simulations predict stable SS/HH_k_-trimers for EGFR^InsNPG^, the FLImP decomposition for this variant reveals no component in the 7–9 nm region (Fig. 3c). This discrepancy likely arises because EGFR^InsNPG^ forms small oligomers (Fig. 3f). This indicates that the back-to-back ectodomain dimer linked to a head-to-head kinase dimer (BB_e_–HH_k_-dimer)^11^ does not readily engage in oligomer assembly for this variant (Fig. 1a). Acting as a structural cantilever, these BB_e_–HH_k_-dimers support the conformationally flexible head-to-head ectodomain (HH_e_) dimers. This support provides the essential robustness needed for oligomers to grow beyond a tetramer assembled by two HH_e_-dimers (Fig. 1c). Furthermore, because BB_e_–HH_k_-dimers provide the essential scaffold for the SS/HH_k_-trimer (Fig. 3b), their absence prevents SS/HH_k_-trimers form forming. This explains the absence of the 7–9 nm peak for EGFR^InsNPG^ (Fig. 3c). Similarly, ED/RK did not significantly reduce the 7–9 nm separations for EGFR^L858R^ (Fig. 3e), which assembles the shortest oligomers in this cohort (Fig. 3f). This result indicates that BB_e_–HH_k_-dimers do not readily engage in EGFR^L858R^ oligomers either. This lack of engagement is unsurprising given the propensity of the L858R mutation to promote the active kinase conformation^14^, and it directly explains the absence of SS/HH_k_-trimers. Ultimately, this validates the assignment of the broad peak at ~10 nm in the EGFR^L858R^ decomposition to separations uncoupled from kinase interactions (Fig. 3b, black arrow, Fig. 3c).

In longer oligomers containing both HH_e_-dimers and BB_e_–HH_k_-dimers, detecting the 7–9 nm separations associated with the SS/HH_k_-trimer requires probes to bind to both dimer species (blue lines, Fig. 3b). For EGFR^L/T^ and EGFR^L/T/C^, the presence of components spanning ~5-6.5 nm (HH_e_-dimer) and ~12-13 nm (BB_e_– HH_k_-dimer) confirm probe binding to these dimers (Figs. 3a, 3c). This guarantees that SS/HH_k_-trimers would be detected if present (Fig. 3b). Consequently, the absence of an SSI-inhibitory effect by the ED/RK mutations reveals that EGFR^L/T^ and EGFR^L/T/C^ do not assemble stable SS/HH_k_-trimers, which agrees with simulations (Figs. 3e, 2b). In contrast, EGFR^T790M^ does not display a component spanning the 12–13 nm range (Fig. 3c). Because EGFR^T790M^ assembles the longest oligomers (Fig. 3f), driven by its strong BB_e_–HH_k_-dimer^15^, the absence of a ~12–13 nm component indicates that most binding sites on these dimers are sterically hindered. Supporting this hypothesis, the ~12-13 nm component reappears when the EGFR^T790M^ oligomers are mutationally disrupted. This recovery occurs without increasing oligomer length, ruling out the recruitment of additional BB_e_–HH_k_-dimers (Extended Data Fig. 5a). Consequently, the absence of an ED/RK effect in EGFR^T790M^ in Fig. 3e is equivocal, as restricted probe binding to the BB_e_–HH_k_-dimer may have prevented the detection of a significant fraction of SS/HH_k_-trimer-dependent separations (blue parallelogram, Fig. 3b). We confirm later that EGFR^T790M^ forms SS/HH_k_-trimers.

The 7–9 nm range also encompasses the ~7 nm separations characteristic of stalk-to-stalk ectodomain dimers coupled to Asym_k_-dimers (StSt_e_–Asym_k_-dimers) (Figs. 1b, 1c). These complexes are unstable as standalone dimers^24,25^. However, as previously shown^11^, they are stabilized within larger oligomeric assemblies through interactions between the BB_e_–HH_k_-dimer and the HH_e_-dimer, the latter of which is linked to two kinase monomers (Fig. 1a). We previously showed that separations around ~7 nm decrease with the TKI lapatinib^11^, a known disruptor of the Asym_k_-dimer^26^. We inferred the presence of StSt_e_–Asym_k_-dimers from the remaining ~7-9 nm separations after introducing the ED/RK mutations designed to disrupt SS/HH_k_-trimers (Fig. 3e). These dimers account for the Asym_k_-dimer-dependent receptor phosphorylation^12,13^ observed in the presence and absence of unnatural mutations (KE and ED/RK) (Fig. 3g).

FLImP data show an anticorrelation between the ~7 nm (StSt_e_–Asym_k_-dimers) and ~9-11 nm (BbBb/Asym_k_-trimers) components (Fig. 3c, yellow and purple components, respectively). Both EGFR^ΔELREA^ and EGFR^L858R^, which do not form BbBb/Asym_k_-trimers, show clear ~7 nm components that are consistent with their increased phosphorylation compared with EGFR^WT^ (Figs. 3c, 3g). While the strong Asym_k_-dimer of EGFR^L858R^ is consistent with previous work^14^, the dimerization strength of EGFR^ΔELREA^ remains debated^27^. Our FLImP data suggest that the EGFR^ΔELREA^ Asym_k_-dimer strength matches that of EGFR^L858R^ (Fig. 3c) aligning with previous experimental data^13^ and MD simulations^14^. EGFR^WT^ similarly fails to form BbBb/Asym_k_-trimers. Consistent with its low phosphorylation (Fig. 3g), a split component in the EGFR^WT^ decomposition spans separations centered at ~7 nm (pale green, Fig. 3c).

Conversely, variants forming stable BbBb/Asym_k_-trimers lack components spanning ~7 nm (Fig. 3c). A clue was provided by the observation that mutational disruption of the EGFR^InsNPG^ oligomer markedly increased the number of ~7 nm separations in this variant (Fig. 3e), in line with the increased levels of StSt_e_-Asym_k_-dimers needed to explain this variant’s hyperphosphorylation (Fig. 3g). MD simulations indicated that these ~7 nm separations can go undetected due to steric hindrance, which can prevent simultaneous binding of two probes to certain HH_e_-dimer conformations (Supplementary Fig. 4), a prerequisite for StSt_e_–Asym_k_-dimer detection (Fig. 3b, orange line). Across the plasma membrane, each HH_e_-dimer links to two kinase monomers, which can be recruited into neighboring kinase dimers to form trimers (Fig. 1). To explain why ~7 nm components go experimentally undetected when the 9–11 nm peaks are prominent (Fig. 3c), we speculate that BbBb/Asym_k_-trimers must induce a conformational change preventing probe binding to one ectodomain of the associated StSt_e_–Asym_k_-dimers (black dot, Fig. 3b).

### Variant-specific structural perturbations dictate ligand-free oligomer assembly rules

Based on assigned components at ~8 nm and ~13 nm and a relatively short oligomer length (Figs. 3c, 3f), we propose that EGFR^WT^ predominantly assembles into hexamers comprising one BB_e_–HH_k_-dimer and two HH_e_-dimers, stabilized by SS/HH_k_-trimers (Fig. 3h). We also propose the following variant-specific differences.

EGFR^InsNPG^ predominantly forms a tetramer of two HH_e_-dimers in which three of the four kinases assemble a BbBb-Asym_k_-trimer (Fig. 3h). This is corroborated by the assigned ~9-11 nm peak and the short oligomer length (Fig. 3c, 3f). The longer tetramer diagonal, measuring ~17 nm (dashed line), falls outside the zoomed-in FLImP decomposition but is prominent in the wider FLImP datasets of EGFR^InsNPG^ and the other variants (Extended Data Fig. 4).

For EGFR^ΔELREA^, we propose two EGFR^WT^-like hexamer motifs linked by an Asym_k_-dimer (Fig. 3h). The assigned components spanning ~7 nm and ~7-9 nm, along with increased oligomer length, support this model (Fig. 3c, 3f). While BB_e_–HH_k_-dimer separations (spanning ~12-13 nm) were undetected in native EGFR^ΔELREA^ (Fig. 3c), they appeared when oligomers were mutationally disrupted and occur without increasing oligomer length (+ED/RK), suggesting initial probe decoration was blocked by steric crowding (Extended Data Fig. 5a).

Short EGFR^L858R^ oligomers alongside the clear component around ~7 nm and the absence of a KE-dependent population of BbBb/Asym_k_-trimers (Figs. 3c, 3d), indicate that EGFR^L858R^ primarily forms tetramers of two HH_e_-dimers held together by a strong Asym_k_-dimer (Fig. 3h). The visible tail in Fig. 3c (green) originates from the ~17 nm peak of the longer tetramer diagonal (Extended Data Fig. 4).

Long EGFR^T790M^, EGFR^L/T^, and EGFR^L/T/C^ oligomers alongside assigned components spanning ~5-6.5 nm and ~9-11 nm (Figs. 3c, 3f) indicate that these oligomers are built upon EGFR^InsNPG^-like tetramer motifs, each containing a BbBb/Asym_k_-trimer, motifs which are cantilevered together by BB_e_–HH_k_-dimers to form the extended oligomers (Fig. 3h).

### Aggressive tumor growth is hardwired to active kinase trimers

The dependence of secondary resistant variants on the BbBb/Asym_k_-trimer may be intrinsic to their oncogenicity. Consistently, our previous work demonstrated that tumors driven by EGFR^T790M^ proliferate more rapidly than those driven by EGFR^T790M+KE^ in the interleukin-3 (IL-3)-dependent Ba/F3 cell system^11^. To reproduce these findings, we generated new Ba/F3 cells stably expressing EGFR^T790M^ (Ba/F3+T790M) or EGFR^T790M+KE^ (Ba/F3+T790M+KE). Our results confirm the attenuated growth of the EGFR^T790M+KE^ variant and reveal a distinct lag phase preceding the onset of exponential proliferation (Fig. 4a).

Within oligomers, StSt_e_-Asym_k_-dimers harbor the high-affinity, growth factor-binding sites essential for driving tumor growth^11,28^. To explain the slower growth of Ba/F3+T790M+KE tumors (Fig. 4a), we previously hypothesized that the integrity of these StSt_e_-Asym_k_-dimers depends on the BbBbI^11^, implying that these high-affinity sites would be disrupted in its absence. However, our MD simulations in Fig. 2b predicted that the BbBbI monomer is dispensable for maintaining the Asym_k_-dimer once formed (Fig. 2b). We experimentally confirmed that StSt_e_-Asym_k_-dimers persist despite inhibition of the BbBbI, showing that both Ba/F3+T790M and Ba/F3-T790M+KE cell lines proliferated at equivalent rates under physiological epidermal growth factor (EGF) ligand concentrations that exclusively bind to these high-affinity sites^28^ (Fig. 4b). Consequently, both our experimental data and simulations implicate the loss of BbBb/Asym_k_-trimers, rather than the loss of high-affinity binding, as the root cause of the slower proliferation observed in EGFR^T790M+KE^ tumors (Fig. 4a).

The proposed loss of BbBb/Asym_k_-trimers is clearly reflected in the FLImP data. As expected, the EGFR^T790M+KE^ variant lacks the 9–11 nm component corresponding to BbBb/Asym_k_-trimers, which are disrupted by the KE mutation. Furthermore, unlike EGFR^T790M^ (Fig. 3c), this double mutant exhibits a prominent component spanning ~12-13 nm, indicating that BB_e_–HH_k_-dimers are doubly labeled (Fig. 4c). The ~12-13 nm component suggests that SS/HH_k_-trimers, if present, should also be detected, consistent with our earlier rationale (blue lines, Fig. 3b). Consequently, the component in the EGFR^T790M+KE^ decomposition centered at ~7 nm could potentially contain signatures from SS/HH_k_-trimers (~7–9 nm), StSt_e_-Asym_k_-dimers (~7 nm), or both (Fig. 4c). To investigate the contribution of the SS/HH_k_-trimers, we added the SSI-disrupting ED/RK mutation to EGFR^T790M+KE^. The resulting FLImP decomposition plot for EGFR^T790M+KE+ED/RK^ (Fig. 4d) recapitulated the results observed for EGFR^T790M^ (Fig. 3c). Specifically, an ED/RK-dependent decrease in the ~7 nm component of the triple-mutant decomposition confirms that EGFR^T790M+KE^ oligomers indeed contained SS/HH_k_-trimers (Figs. 4c, 4d). Concurrently, the ED/RK-dependent restoration of the peak spanning ~9–11 nm shows that a reduction in SS/HH_k_-trimers has shifted the equilibrium back towards BbBb/Asym_k_-trimers, as previously proposed^11^ (Fig. 4d). As anticipated, the reappearance of BbBb/Asym_k_-trimers in EGFR^T790M+KE+ED/RK^ correlates with the absence of a ~7 nm StSt_e_-Asym_k_-dimer component (Fig. 4d). This finding aligns with our prior hypothesis that FLImP cannot detect these two species simultaneously (Fig. 3c).

To determine whether re-establishing BbBb/Asym_k_-trimers impacted tumor growth, we generated Ba/F3 cells stably expressing the quadruple mutant EGFR^T790M+KE+ED/RK^ (Ba/F3+T790M+KE+ED/RK). Unlike tumors derived from Ba/F3+T790M+KE cells, those derived from these quadruple-mutant cells lack the distinct lag phase and achieved growth rates closer to Ba/F3+T790M tumors (Fig. 4a, Extended Data Table 1a). These findings demonstrate that the BbBb/Asym_k_-trimers are crucial mediators of EGFR^T790M^-driven tumor growth.

### Catalytic kinase trimers rewire downstream signaling

To understand how these BbBb/Asym_k_-trimers promote accelerated tumor growth, we investigated how their conversion into Asym_k_-dimers influences receptor behavior and downstream pathways. At the plasma membrane, the different entropic costs and degrees of freedom of trimers versus dimers are reflected by changes in oligomer assembly kinetics and diffusion rates (Extended Data Fig. 5b). Furthermore, trimers and dimers can either expose or obscure distinct adaptor- or effector-binding sites, thereby modulating downstream signalling. This was revealed by a targeted antibody array profile of IL-3-starved Ba/F3+WT, Ba/F3+T790M, and Ba/F3+T790M+KE cells, using Ba/F3+T790M as a control (Fig. 5a). Analysis of significantly altered proteins (identified by Z score^29^) showed that the KE mutation led to a 33% decrease in the expression of mouse double minute 2 (MDM2), an E3 ubiquitin ligase and master negative regulator of the tumor suppressor p53 transcription factor, and a 34% increase p53 phosphorylation at S15^30^. Phosphorylation at S15 activates p53, which compromises the ability of the depleted MDM2 pool to bind and target it for proteasomal degradation^31^.

**Figure 5.**
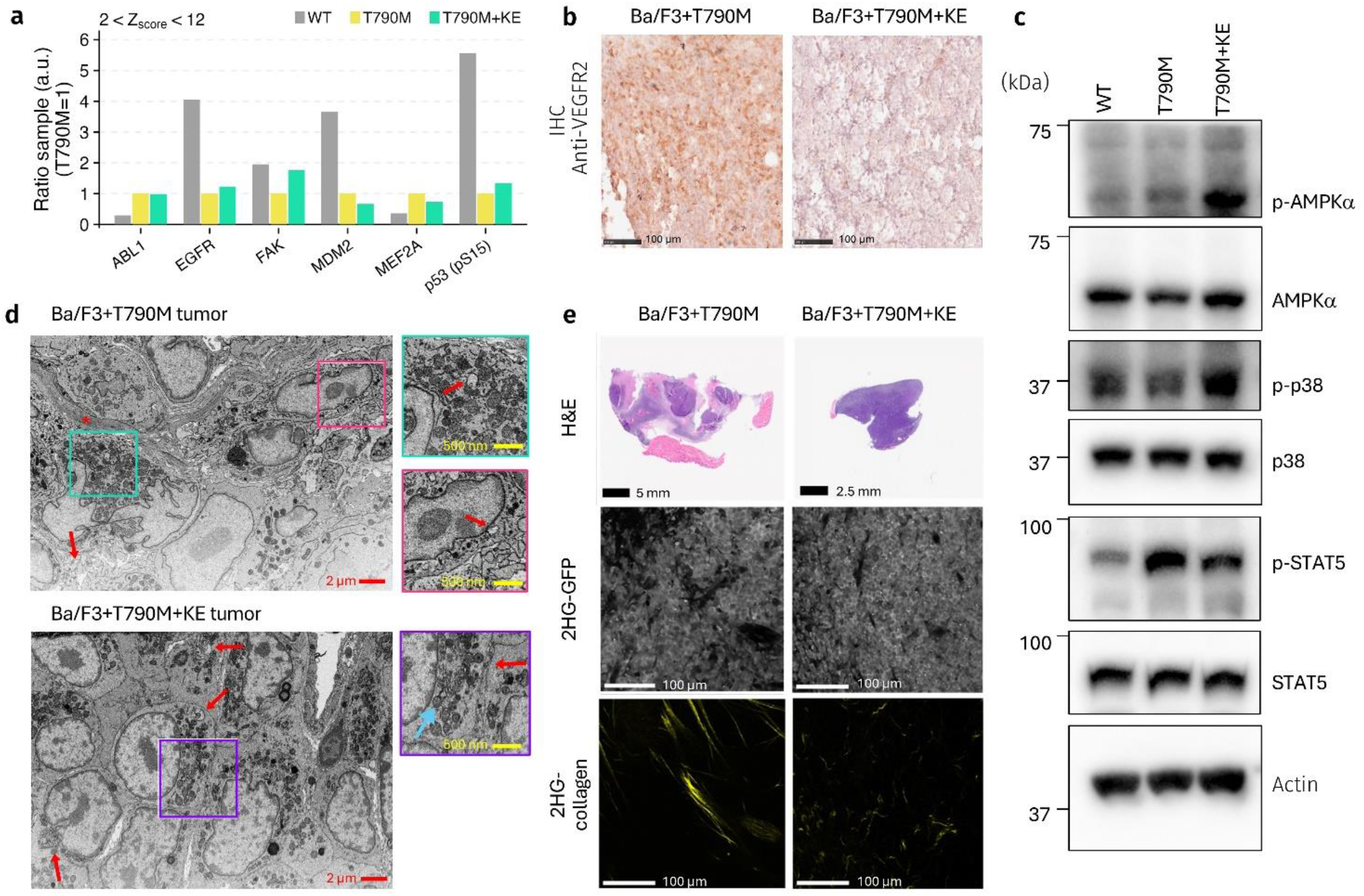
The trimer-to-dimer transition rewires downstream signaling. **a**, Quantitative profiling of protein expression and phosphorylation derived from a targeted protein antibody array, showing results with a Z-score^29^ of 2-12 (Details in Extended data Fig. 9). **b**, Immunohistochemistry staining of Ba/F3+T790M and Ba/F3+T790M+KE tumor sections using an antibody against VEGFR2 (See also Extended data Fig. 8b). **c**, Western blot showing phosphorylation of AMPKα, p38, and STAT5 in the absence of growth factor in stably transfected Ba/F3 cells. Blot quantification is shown in Extended data Fig. 6c. **d**, Representative volume EM images of Ba/F3+T790M (top) and Ba/F3+T790M+KE (bottom) mouse tumors. Scale bars, 2 µm (main panels) and 500 nm (insets). Boxes indicate the inset regions. Top: Red arrows indicate regions of interdigitated cell-to-cell contacts; asterisk indicates a bundle of collagen fibers. Inset arrows point to late endosomes/lysosomes. Inset 1: close-up of mitochondria showing dense arrangement of cristae. Inset 2: close-up of interdigitated membrane contact area. Bottom: arrows indicate enlarged late endosomes/lysosomes. Inset: close-up of mitochondria showing hole-like defects (cyan arrow). Red arrows indicate close, tight cell-to-cell contacts. **e**, Representative histology and light microscopy images of Ba/F3+T790M and Ba/F3+T790M+KE mouse tumors (top). Top: Haematoxylin and eosin (H&E) images collected on a slide scanner. Scale bars, 5 mm (Ba/F3+T790M) and 2.5 mm (Ba/F3+T790M+KE). Middle: second-harmonic generation images of GFP-tagged tumor cells. Scale bar, 100 µm. Bottom: second-harmonic generation images of unstained collagen. Scale bar, 100 µm.

Depending on cellular p53 concentration, the KE-dependent accumulation of stabilized p53 can either trigger apoptosis or drive cell cycle arrest at G1/S and G2/M checkpoints^32^. This cell cycle arrest likely explains the distinct lag phase of Ba/F3+T790M+KE tumors (Fig. 4a). An increase in p53 can also downregulate angiogenesis by inhibiting pro-angiogenic effectors such as vascular endothelial growth factor and its receptor (VEGFR2), as found in glioma^33^. Consistent with this anti-angiogenic role, we found that the KE mutation downregulated VEGFR2 in the Ba/F3+T790M+KE tumors (Fig. 5b). Upstream of the MDM2/p53-pS15 axis, stress-response pathways were activated, as evidenced by the KE-dependent increased phosphorylation of AMP-activated Protein Kinase alpha (AMPKα) and its downstream effector p38 (Fig. 5c). AMPKα regulates the glucose-dependent checkpoint at the G1/S boundary via p53 phosphorylation at S15^34^ whereas p38, a master regulator of inflammation and stress responses, promotes MDM2 degradation^35^. Complementing these changes, the KE mutation simultaneously suppressed oncogenic survival signaling, as demonstrated by decreased phosphorylation of Signal Transducer and Activator of Transcription 5 (STAT5) a key effector through which oncogenic EGFR rescues cells from apoptosis induced by IL-3 depletion^36^ (Fig. 5c).

The KE mutation also repressed the expression of Myocyte Enhancer Factor 2A (MEF2A) by 26% (Fig. 5a). MEF2A serves as a major transcriptional coordinator of mitochondrial homeostasis during cellular stress^37^. Because MEF2A downregulation would reduce mitochondrial metabolic support for tumor growth, we imaged mitochondria in tumor sections using volume electron microscopy. Images showed mitochondrial damage, consistent with a MEF2A-dependent decrease in mitochondrial biogenesis, alongside an increase in lysosomes, both reflecting metabolic stress^37,38^ (Fig. 5d).

The KE mutation also induced a 76% increase in focal adhesion kinase (FAK) expression^39^ (Fig. 5a). As a non-receptor tyrosine kinase, FAK not only regulates cell adhesion, migration, and survival but also remodels the extracellular matrix and tumor microenvironment^40^. Consistent with the former, Ba/F3+T790M+KE tumors are denser than Ba/F3+T790M ones (Fig. 5e, top). Consistent with the latter, second-harmonic generation (SHG) microscopy revealed the disruption of tumor-associated interstitial collagen fibers (Fig. 5e, bottom). Together, results in in Fig. 5 demonstrate that the KE mutation acts as a potent metabolic stressor and an oncogenic suppressor.

### The trimer-to-dimer transition re-sensitizes tumors to gefitinib

Consistent with its known role in proliferation, the expression of the non-receptor tyrosine kinase Abelson murine leukemia viral oncogene homolog 1 (ABL1)^41^ increased fourfold in both Ba/F3+T790M and Ba/F3+T790M+KE relative to Ba/F3+WT cells (Fig. 5a). This reveals that ABL1 expression—sustained by FAK— maintains tumor proliferation and survival in the Ba/F3+T790M+KE model.

ABL1 binds to and phosphorylates EGFR, shielding EGFR from ubiquitination and proteasomal degradation by inhibiting the accumulation of the E3 ubiquitin ligase Cbl at the plasma membrane^42^. ABL1 is also a downstream target of EGFR^43^. We hypothesized that a conformational change in EGFR^T790M+KE^ might disrupt downstream EGFR–ABL1 signaling. To test this, we treated Ba/F3+T790M+KE cells with 1 µM gefitinib, a concentration more than 90-fold higher than the dissociation constant (*K*_*d*_) of gefitinib binding to the ATP-pocket of the EGFR^T790M^ kinase domain^44^. Gefitinib competes with ATP for binding to the ATP-binding pocket of EGFR’s kinase domain and is known to alter EGFR conformation^26^. This was reflected by a conformational change in EGFR^T790M+KE^ oligomers (Extended Data Fig. 5c). Compared to the lowered ATP-binding binding affinity of gefitinib-sensitive classical mutations^45,46^, the T790M mutation increases ATP-binding affinity by ~25-fold to ~6 µM, almost restoring it to EGFR^WT^ levels^44^. Unsurprisingly, given the ~5 mM ATP concentration in mammalian cells^47^, EGFR^T790M+KE^ phosphorylation remained unchanged in the presence of gefitinib (Extended Data Fig. 6b); however, gefitinib treatment downregulated ABL1 expression four-fold (Fig. 6a). This suggests that the gefitinib-induced conformational change in EGFR^T790M+KE^, rather than a loss of kinase activity, is the primary driver of the reduced ABL1 stability.

**Figure 6.**
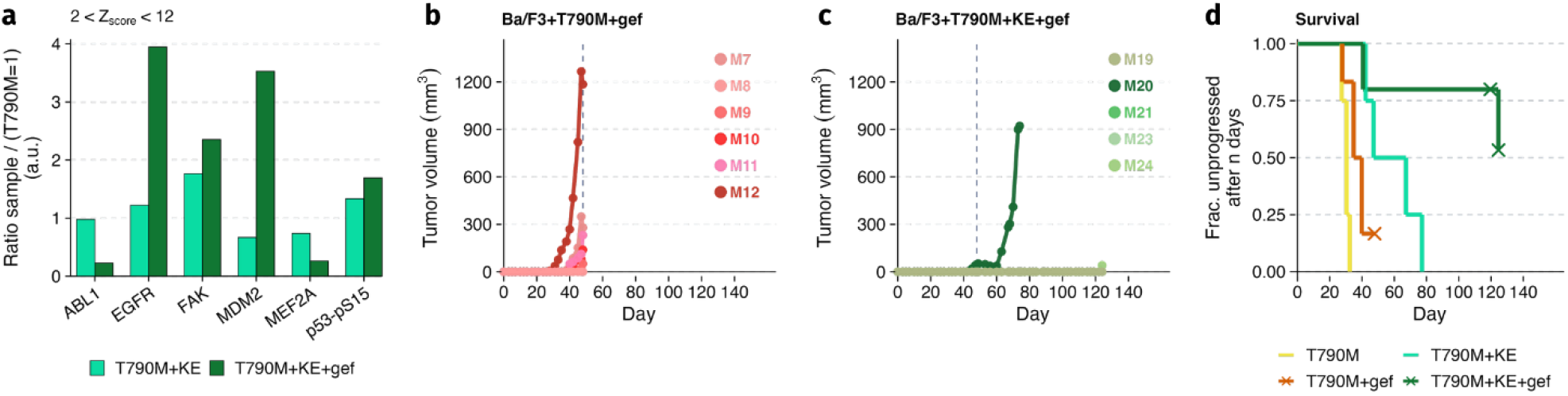
A trimer-to-dimer transition overcomes T790M-mediated gefitinib resistance. **a**, Quantitative profiling of protein expression and phosphorylation derived from a protein antibody array. Displayed targets met a threshold of a Z score^29^ = 2–12 (Extended data Fig. 9). **b, c**, *In vivo* tumor growth kinetics in NSG mice bearing subcutaneous xenografts of Ba/F3 cells expressing EGFR^T790M^ or EGFR^T790M+KE^ treated with gefitinib. Tumor volume was measured longitudinally using calipers. *n* = 6 biologically independent animals per cohort; one animal was censored in (c). The vertical dashed line indicates day 48, when the T790M cohort reached the end point and was culled (Details in Extended data Fig. 7). **d**, Kaplan-Meir survival plot^51^. Crosses denote censored mice.

Despite the downregulation of ABL1, which binds p53 and neutralizes the inhibitory effect of MDM2^48^, the level of activated p53-pS15 increased further with gefitinib (Fig. 6a). This p53 activation accounts for the transcription upregulation of MDM2 (Fig. 6a). Coupled with the four-fold increase in the expression EGFR^T790M+KE^, these findings suggest a gefitinib-induced cell-cycle arrest or senescence driven by parallel p53-dependent and oncogenic-induced pathways^48,49^. Consistent with this, murine model evaluations demonstrated that while gefitinib failed to suppress the growth of Ba/F3+T790M tumors, it successfully suppressed the growth of Ba/F3+T790M+KE tumors (Figs. 6b, 6c). These results mirror the clinical response rates to gefitinib observed in NSCLC patients harboring the classical, gefitinib-responsive ΔELREA and L858R mutations (80.3% and 81.8% respectively)^50^.

Kaplan-Meir survival analysis^51^ revealed a modest difference between the EGFR^T790M^ plus or minus gefitinib cohorts (p=0.0400), which became markedly more significant when comparing the EGFR^T790M^ and EGFR^T790M+KE^ + gefitinib cohorts (p=0.0029) (Fig. 6d, Extended Data Table 1b). An independent analysis of the tumor growth trajectories between treatment systems at day 128 showed significant differences between the EGFR^T790M+KE^ plus and minus gefitinib cohorts (p<0.0001) (Extended Data Table 1b). These results show that gefitinib significantly inhibits the growth of tumors driven by EGFR^T790M+KE^, thereby demonstrating that trimer-to-dimer transition renders the corresponding tumors sensitive to gefitinib treatment. While our complete functional characterization focused on the EGFR^T790M^ background, our MD simulations and structural data across variants suggest a shared dependence on BbBb-Asym_k_-trimer integrity. Combined with the effect of the BbBb-Asym_k_-trimer disruption and gefitinib on the endocytosis of EGFR^L/T/C^ (Supplementary Fig. 5), these insights firmly establish catalytic kinase trimers as drivers of acquired resistance and their disruption as a generalizable therapeutic strategy to overcome this resistance.

## Discussion

The concept that receptor kinases may reversibly assemble catalytic units larger than dimers was suggested a decade ago^52^. However, this has remained hypothetical, including in our own earlier work^11^. To address this gap, in this study, we demonstrate that two generations of acquired TKI resistance are underpinned by a transition from canonical active dimers, characteristic of EGFR^WT^ and classical, TKI-permissive mutants^12,14^, to unconventional catalytic trimers. Within these trimers, an ancillary BbBbI overcomes the intrinsic instability of the Asym_k_-dimer^13^. Crucially we show that disrupting these unconventional trimers and restoring canonical dimers via a single-point genetic rescue impairs tumor growth and re-sensitizes the T790M mutation to gefitinib.

To bridge oligomer architecture with *in vivo* oncogenic phenotypes, we deployed a multi-scale interdisciplinary framework. First MD simulations of the non-canonical active and inactive trimers were validated on cells via ~2 nm-resolution FLImP microscopy. Next, microarray profiling of downstream signaling, confirmed by Western blotting and volume EM and SHG imaging of tumor sections, revealed how disrupting the unconventional catalytic trimers fundamentally alters the EGFR signaling network, thereby inhibiting tumor growth and resensitizing T790M to gefitinib. Thus, these trimers are critical, targetable drivers of clinical relevance.

Consequently, targeting these trimer lesions in EGFR-mutant oligomers offers an untapped opportunity for structure-based drug design, laying the foundations for future strategies to combat drug resistance. By uncovering a conformational state unique to acquired-resistance-associated oncogenes, targeting the kinase trimer-to-dimer transition provides a mutant-selective therapeutic window. This approach circumvents the wild-type toxicities associated with non-specific extracellular oligomer disruption^10,53^ and Asym_k_-dimer inhibition^54^. Additionally, it avoids the complex toxicity profiles of standard combination therapies that target parallel survival pathways^55^.

Identifying the ancillary interfaces that regulate active and autoinhibited trimer–dimer transitions aligns with rapid advances in protein–protein interaction modulators^56^. Reverting active trimers to dimers via a single-point mutation maps a structural ‘hotspot’ within the BbBbI^57^, simplifying the future design of targeted minibinder inhibitors^58^. Delivering these minibinders via mRNA platforms^59^ alongside first-generation TKIs could durably suppress acquired resistance periods, because successive generations of TKI-resistant variants share these catalytic trimer lesions. Ultimately, targeting the ancillary BbBbI could represent a universal strategy against secondary TKI resistance that could transform the clinical efficacy of existing TKIs, providing cost-effective therapeutic regimens that extend patient disease-free survival with fewer side effects.

## METHODS

### Reagents

A list of reagents can be found in Supplementary Information.

### Cell culture

Chinese Hamster Ovary (CHO) cells (gift from Prof. Peter Parker at The Francis Crick Institute, UK) and Ba/F3 cells (Creative Biogene, CSC-C2045) were grown according to standard protocols and as described previously^11^. Additionally, they were verified negative for mycoplasma before use and tested routinely.

### Plasmid construction

Point mutations in EGFR plasmids were generated by site-directed mutagenesis using either the QuikChange Lightning kit (Agilent Technologies) for EGFR/pcDNA3 constructs or the Q5 kit (NEB) for EGFR/PB513B-1 constructs, with primers listed in Supplementary Table 2. All constructs were validated by sequencing the complete EGFR coding sequence.

### Generation of Ba/F3 or CHO stable cell lines

Ba/F3 and CHO cells were electroporated using a Neon transfection kit (Invitrogen, MPK10096) and MicroPorator device as before^11^. Stably expressing cells were maintained in 2 µg/µL or 4µg/µL puromycin dihydrochloride, respectively.

### FAC sorting

Cells were washed twice in ice-cold PBS containing 5% FBS and 2 mM EDTA and sorted under sterile conditions using a BD FACS Melody. GFP thresholds were set based on the highest GFP signal of the lowest-expressing population and applied across all populations to isolate singlet cells with matched GFP, and hence EGFR, expression. Cells were recovered in complete medium containing puromycin and IL-3. All sorted cell lines were confirmed mycoplasma-negative.

### Ba/F3 IL3 independent growth assay

Ba/F3 cells stably expressing T790M-EGFR or T790M+KE-EGFR were washed to remove IL-3 and puromycin and cultured for 5 days. Cells (20,000 per well) were then seeded in triplicate in white 96-well plates containing 100 μl medium supplemented with 200 pM EGF. Cell viability was monitored for 12 days using CellTiter-Glo (Promega, G7572) and a CLARIOstar Plus microplate reader (BMG Labtech), according to the manufacturer’s instructions.

### Microarray data collection and analysis

Ba/F3 cells expressing WT-EGFR, T790M-EGFR, T790M+KE-EGFR or T790M+KE+ED/RK were washed once with PBS and cultured without IL-3 nor puromycin for 5 days, followed by 24 h in medium containing 100 pM EGF. Where indicated, cells were treated with 1 μM gefitinib (Sigma-Aldrich, SML1657) or DMSO vehicle for 1 h. Cells were washed three times with ice-cold PBS and lysed in RIPA buffer (Thermo Scientific, 89900) supplemented with 1 mM dithiothreitol, 1 mM sodium orthovanadate and protease inhibitors (Cell Signaling Technology, 5871). Lysates were incubated on ice for 50 min with vigorous vortexing every 10 min, clarified by centrifugation and quantified by Bradford assay.

A microarray specific to EGFR-signaling pathways (Creative Biolabs, #Z01MM-0326-M1) detected changes in phosphorylation and protein levels relative to the T790M-EGFR control sample using 239 specialized antibodies (the protocol, list of antibodies and raw data are available on Zenodo at https://doi.org/10.5281/zenodo.22128078). For each antibody, signal was normalized using the “F532_median” and “F532_median” readouts as foreground and background, respectively and the signal of multiple spots was averaged. Z-Scores were calculated as z_i_ = (x_i_ – <x>/σ_x_), as described in^29^. Fold change was calculated per gene against expression in the T790M dataset. Proteins were considered hits if Z-score >1 (expression > 1σ above assay average) and fold change > 2 (upregulated) or <0.5 (downregulated). Data was plotted using Excel or R with ggplot library.

### Western blot

CHO cells at approximately 25% confluency were transfected with 2 µs of the appropriate WT or mutant EGFR DNA using Jet Optimus (Sartorius #101000051) at a ratio of 1µg DNA:1µL reagent according to the manufacturer’s instructions. 24 h post-transfection, the cells were serum starved in media containing 0.1% FBS for >16 h. Cells were placed on ice, washed in cold PBS, lysed and clarified by centrifugation as described previously^11^. 5-7.5 µgs of total cell lysate protein, as determined using the Bradford assay, was resolved on a 4-12% Bis-Tris gel (NuPAGE #WBT41226), and proteins were transferred to PVDF membrane. The membranes were blocked, incubated in primary and secondary antibodies according to the manufacturers recommended dilutions and as described previously^11^. Subsequently, membranes were incubated with Immobilon (Millipore WBKLS0500) or Pierce ECL Ultra Western (Pierce # 32209) HRP substrate solutions and imaged using a Biorad ChemiDoc™ XRS+ molecular imager. The images were quantified using Image Lab™ 6.1 software where required. Alternatively, 15 μgs of total protein from the BA/F3 cell lysates from the microarray samples detailed above were handled in the same way.

### FLImP Sample Affibody labeling

CHO cells expressing EGFR^KE^, EGFR^insNPG^, EGFR^insNPG+KE^, EGFR^insNPG+ED/RK^, EGFR^ΔELREA^, EGFR^ΔELREA+KE^, EGFR^ΔELREA+ED/RK^, EGFR^L858R+KE^, EGFR^L858R+ED/RK^, EGFR^L/T^, EGFR^L/T+KE^, EGFR^L/T+ED/RK^, EGFR^L/T/C^, EGFR^L/T/C+KE^ and EGFR^L/T/C+ED/RK^ by Jet optimus transfection (300 ng plasmid DNA, at 1:1 DNA:Jet Optimus ratio Sartorius #101000051) or stable cell lines for EGFR^WT^, EGFR^ED/RK^, EGFR^L858R^, EGFR^T790M^, EGFR^T790M+KE^ and EGFR^T790M+ED/RK^ were grown to <80% confluency in 1% BSA-coated wells of μ-Slide, 8 well, high glass bottom slides (ibidi, 80807).The 4th well (top left) was coated with poly-L-Lysine (PLL) only. Samples were incubated in low serum medium (0.1% FBS), with 1 µM gefitinib if necessary, for 2 h before labelling with 8 nM HER1 Affibody-CF640R made in house, thorough fixing and labeling with1 µg/mL Hoechst 33258 (Life Technologies, #H3569) as described previously^11^. Fiducial markers (1/200,000 dilution of a 2% solution of FluoSpheres™, 0.1 µm, infrared (715/755) (Invitrogen F8799 in PBS) were added to all wells and samples were loaded onto the microscope as described previously^11^.

### FLImP data acquisition

Image acquisition was achieved using an Oxford Nanoimager S (ONI Oxford, UK) single molecule imaging microscope with a 1.49N oil immersion objective, operating NanoImager software (Version: 1.7.3.10248 – ef4ff2c0) set up according to the manufacturer’s instructions. For more details see^11^.

### Animals

In this study, young adult male (6 weeks old, 24.36 ± 1.91 g) NOD.Cg-Prkdc^scid^ Il2rg^tm1Wjl^/SzJ mice (NSG; purchased from Charles River UK) were used for all animal experiments. All mice were maintained within the King’s College London Biological Services Unit as described previously^11^. All experiments were approved by the King’s College London Animal Welfare and Ethical Review Body in accordance with UK Home Office regulations (Project License PP4067431) and UK NCRI Guidelines for the Welfare and Use of Animals in Cancer Research. Experimental design was based on the assumption that tumor growth differences *in vivo* are comparable to the effects observed upon Gefitinib/Iressa administration. This data was used together with an α of 0.05 and a power of ≥90% to determine minimum cohort sizes. For longitudinal experiments, cohort sizes were then oversubscribed to hedge against potential adverse effects and resultant animal sacrifice, which, if premature, would endanger the whole study. Consequently, cohort sizes were *N*=6. The total number of animals used was 30. No adverse events were associated with the procedures performed in this study and animals put on weight in line with strain expectations (data from Charles River UK) throughout. Sentinel animals were kept on the same IVC racks as experimental animals and confirmed to be healthy after completion of the studies.

### Tumor models

Male NSG mice were used to establish subcutaneous tumor models (in right flanks) with indicated stable Ba/F3 cell lines. After acclimatisation, mice were randomised into five cohorts with six individuals each, shaved on their flanks, and then subcutaneously received each 2×10^6^ tumor cells suspended in 100 μL phosphate buffered saline (PBS). Starting from Day 0, the mice either received intraperitoneal injection of Gefitinib (Iressa, Tocris Bioscience, #3000) at 40 mg/kg or an equivalent dose of vehicle (DMSO) thrice a week. Tumor growth was followed by callipers and tumor volumes were calculated using the formula: 0.5 × L × W^2^, wherein L represents tumor length and W its width. Tumor models were grown to compare tumor growth between cohorts. The experimental endpoint was determined by the time the human endpoint was reached, where a single tumour reached a Maximum Tumor Diameter (MTD) of 1.2 cm. *In vivo* GFP fluorescence imaging of superficial tumor models was performed to visualise tumor growth in some animals per group over time and to quantify tumour growth differences in all animals at the experimental endpoint^11^.

### Tissue staining and histologic tissue analysis

Formaldehyde-fixed paraffin-embedded (FFPE) tissues were prepared and stained with either haematoxylin- and eosin (H&E) for morphologic analysis or antibodies (Anti-EGFR, D38B1, Cell Signaling Technology (4267) or Anti-VEGFR, D5B1, Cell Signaling Technology 9698 or Anti-GFP at final concentrations of 100ng/mL, 20µg/mL and ~90ng/mL respectively in 2%BSA in TBS) to test for protein expression using standard methods and as described before^11^. Samples were developed using the Pierce DAB Substrate Kit (ThermoFisher, UK). Slides were scanned using a Nanozoomer (Hamamatsu, Japan), with images being analysed and processed by ImageJ v1.54 (NIH, USA).

### Volume EM imaging of mouse tumor samples

Sample dissection and fixation followed the workflow described in^61^ with modifications. Tumor tissue was fixed promptly after dissection for 1 h at RT with 4% PFA (EM Grade, Electron Microscopy Sciences 157-4) in 0.1 M phosphate buffer (PB) and then subsequently stored at 4°C in 1% PFA in 0.1 M PB. Tissue blocks were embedded in 3% agar in 0.1 M PB and cut into 200 μm sections using a 7000 smz-2 vibrating blade tissue slicer (EMS) at 80Hz, amplitude = 2.0 mm, speed = 0.25 mm/sec while immersed in 0.1 M PB buffer. Sections were post-fixed with 4% PFA + 2.5% GA in 0.1 M PB for 30 min, then washed in 0.1 M PB, 5 × 3 min and trimmed to 0.5 mm^2^

Samples were stained and resin embedded at the Oxford Dunn’s School of Pathology EM facility using a modification of the NCIMR protocol^62^. Briefly, samples were stained with reduced osmium (2% OsO_4_, 1.5% K_3_Fe(CN)_6_) for 1 h at 4°C without agitation, then with 1% thiocarbohydrazide (TCH) for 20 min at RT, then with 2% aqueous OsO_4_ in ddH_2_O for 30 min at RT, with each staining step being followed by washing 3 × 5 min in ddH_2_O at RT with agitation. Uranyl acetate staining was performed with 1% uranyl acetate at 4ºC overnight, followed by washing in ddH_2_O, 5 × 3 min at RT and then lead aspartate staining for 30 min at 60°C. Samples were washed again in ddH_2_O, 5 × 3 min at RT, and then dehydrated in ethanol series 30%-50%-70%-90%-100% - dried ethanol, with each step being performed for 10 min. Samples were then washed in acetone for 10 min and then infiltrated with Durcupan in acetone at 25:75 then 50:50 and then 75:25 v/v, with each step taking 2 h. Infiltration in 100% Durcupan was performed overnight at RT, then samples were changed to fresh 100% Durcupan for 2 h, embedded into BEEM® capsules and cured for 48 h at 60°C. Test semi-thin sections (1-3 µm) were stained with 1% Toluidine Blue and imaged under a dissection microscope for Quality Control purposes.

Samples were imaged in a Zeiss Crossbeam 550L over volumes of at least 40 × 30 × 20 μm at 10 nm^3^ isometric voxels. Milling was performed at 30 kV 700 pA, dwell 228 μsec, dose 1600mC/cm^2^. Imaging was performed at 1.5 kV 750 pA, with scan speed 5, with noise reduction using line averaging over 4 lines and a dwell time of 100 nsec. The signal was detected on the in-column ESB detector with the grid set to 800V.

### Second Harmonic Generation (SHG) imaging of mouse tumor samples

200 μm thick sections of lightly fixed (4% PFA in 0.1 M PB) tumor were imaged in a glass-bottom dish in 0.1 M PB while weighed down with a 22 mm round coverslip to limit sample movement during XY stage movement.

SHG Imaging was performed on a Leica SP8 instrument. Images were collected at a pixel size 0.143 μm, using the PMT-RLD 3 detector with filters BP483/32 and gain 500 for GFP, and the PMT-RLD 4 detector with filter BP 440/20 and gain 750 for collagen. Samples were imaged using a Fluotar VISIR 25x/0.95 water immersion objective and illuminated with 4% power at 880 nm from an Insight DeepSee multiphoton laser from Spectraphysics at a scan speed 700 Hz. Pinhole size was 50.1 μm.

### Tumor growth analysis

Tumor growth was analyzed with a linear mixed-effects model fit to log1p-transformed tumor volume, which retains zero-volume observations. Fixed effects were day, treatment system, and their interaction, with mouse-specific random intercepts and random slopes for day to account for within-mouse correlation across repeated measurements. Models were fit by maximum likelihood, and an overall difference in growth trajectories across systems was assessed with a likelihood-ratio test comparing the full model against a reduced model without system terms. Pairwise system contrasts at day 48 and day 124 were evaluated as model-estimated log-differences and fold-change ratios. Full statistics are given in Extended data Table 1 and Supplementary Note 4.

### Confocal imaging of endocytosis samples

CHO cells were transfected with the indicated EGFR constructs using JetOPTIMUS. After 48 h, cells were washed, serum-starved for 2 h, and incubated in serum-free medium with or without 1 μM gefitinib, as indicated. Cells were then chilled on ice (4°C, 10 min) and incubated for 1 h at 4°C with 100 nM CF640R-conjugated anti-EGFR EgB4 nanobody with 1 μM gefitinib if required. Following extensive washing, cells were stimulated with 200 pM EGF, with 1 μM gefitinib if required, for 30 min at 37°C. Cells were immediately fixed in 3.5% paraformaldehyde (EM Grade; EMS 1574) for 30 min at room temperature, stored at 4 °C, and labelled before imaging with 5 μg/ml wheat germ agglutinin–Alexa Fluor 488 (Invitrogen, W11261).Samples were imaged on Zeiss Elyra PS1 using a 63x/NA 1.45 oil immersion objective. WGA-AF488 nm and EgB4-CF640 nm images were acquired sequentially using 0.6% 488 nm and 2% 633 nm laser power and 900 AU and 600 AU gain respectively and the following invariant parameters: pinhole size = 1.35 AU, pixel dwell time = 1.72 μsec, tile size 1220 × 1220 pixel, pixel size 0.11 μm, tilescan acquisition 2×2.Image analysis was performed in FiJi^63^ as follows: for each dataset, channels C0 (EgB4-CF640R) and C1 (WGA) were split then a rolling ball background subtraction of 20 px was applied to C0.A threshold of 50 intensity counts was applied to C1 and the image was dilated and renamed as “mask”, then duplicated and the duplicate mask was inverted and renamed “invert_mask”. The “mask” image was subtracted from C0 using the Image Calculator function and the resulting image was thresholded using a value of 50 intensity counts.Object-based statistics were obtained using the Analyse Particles function (diameter 0.04-1 um, circularity 0.4-1). Objects were added to the ROI Manager and transferred to the background-subtracted C0 image, and the per-object integrated density was measured. Data was saved as XXNN_endo.csv, where XX is the condition code and NN is the ROI number. To measure the same set of parameters for membrane objects, the “invert_mask” image was subtracted from C0 using the Image Calculator function and the resulting image was thresholded using a value of 50 intensity counts, then object-based parameters were measured as above, except that Analyse Particles function was applied with the following parameters: diameter = 0.5-Infinity um, circularity = 0.0-0.4. Results were saved as XXNN_membrane.csv. A custom Python script was used to parse all csv files and compile a dataframe containing the sum of the integrated density of endocytosed and membrane objects per-ROI, as well as the integrated density ratio. Data was plotted in Python using the Seaborn and Matplotlib libraries. Statistics were calculated using the Kruskal-Wallis test implemented in the scipy.stats library and the posthoc_ttest function implemented in the scikit_posthocs library with Bonferroni correction for multiple comparisons. Raw data, dataframes and code are available on Zenodo at: https://doi.org/10.5281/zenodo.22129114

### Single particle tracking

CHO cells were prepared and imaged as described previously^11^. All single-molecule time series data (for FLImP and single particle tracking) were initially analyzed using the multidimensional analysis software described previously^64^. The colocalization event duration analysis was performed in the same way as in^15^.

### FLImP data analysis

Details of FLImP separation decompositions and oligomer length determination can be found in Supplementary Figs. 2 and 3 and was previously described^11^.

### MD simulations of kinase trimers

Dimer models were taken directly from deposited crystal structures. The Asym_k_-dimer (PDB entry 2GS6^12^) and HH_k_-dimer (PDB entry 5CNO^65^) are resolved in the main electron density of their respective PDB entries, while the BbBb_k_-dimer (PDB entry 3VJO^66^) and SS_k_-dimer (PDB entry 5CNO) are found in the crystal lattice of deposited structures. The BbBb/Asym_k_-trimer and SS/HH_k_-trimer are speculative models that we built using the resolved dimeric structures to interpret experimental data, following the modelling and validation strategy described previously^11^. The modeling details for the trimeric structures are given in Supplementary Note 1. Mutations were introduced onto the WT trimeric structures with MODELLER^67^. The full set of mutations and sequence boundaries used for each system are listed in Supplementary Note 1 and Supplementary Table 2. Because our protocol simulates preformed trimers, it does not capture any mutation-driven effects on monomer folding that might prevent dimer or trimer assembly in the first place. However, FLImP separation data indicate that trimers do assemble under these conditions, supporting this modeling assumption.

Each trimeric system was parameterized with the CHARMM36m force field^68^, protonated at pH 7.4 using the Proteinprepare tool of the PlayMolecule suite^69^, solvated in a dodecahedral box of modified TIP3P water^70^, and neutralized with Na^+^/Cl^−^ to 0.15 M. All simulations were carried out using GROMACS v.2022.5^71^. Systems were energy-minimized and equilibrated (NVT then NPT) before unbiased production runs. The full equilibration and production details are given in Supplementary Note 2. Simulation lengths and replica numbers for each system are listed in Supplementary Table 1.

Interface strength was calculated with mdciao^72^, based on residue–residue distance time traces at each dimer interface (4 Å cutoff), and is reported as the average number of interface contacts. Simulations produced one of three outcomes: a numeric strength value, an interface fracture (X), or an interface change (IC), which we report explicitly rather than collapsing into a single average. The rationale behind this choice is given in Supplementary Note 3.

## Acknowledgements

We thank Martin Attwood, Outfox Bio for help with FACS. This work has been funded by grant Ref: ST/Y004183/1 from the Science and Technology Facilities Council UK. M.L.M.-F., D.J.R., and B.M.D. are grateful for IRIS computing resources funded by the Science and Technology Facilities Council (STFC) with operational support from STFC’s Scientific Computing Department. F.L.G., N.P., and I.G. also acknowledge the Swiss National Supercomputing Centre (CSCS) for large supercomputer time allocations, projects s1228, lp84, and lp166.

## Author contributions

Conceptualization, S.K.R., I.G., F.L.G., D.T.C., and M.L.M-F; molecular and cell biology, S.K.R.; MD simulations, IG and NP, who contributed equally to this work; FLImP data collection, S.R.N and B.M.D; PCS data collection and analysis, A.H.A.C.; FLImP data analysis, D.J.R; in vivo work, R.C.H.M., G.O.F., and L.Z.-D.; writing – original draft, M.L.M.-F.; volume EM and SHG imaging, L.Z.-D; writing – review & editing, S.K. R., I.G., N.P., S.R.N., B.M.D., L. Z.-D., R.C.H.M., D.T.C., A.H.A.C., D.J.R., G.O.F., F.L.G.

