## Supplementary figures, tables, and notes for "Disrupting aberrant EGFR catalytic trimers reverses T790M gefitinib resistance"

### CONTENTS:

**Supplementary Fig. 1. Cartoon and ribbon representations of simulated trimers.**

**Supplementary Fig. 2. Key stages in the FLImP data acquisition and analysis process.**

**Supplementary Fig. 3. FLImP measurement-dependent oligomer length determination.**

**Supplementary Fig. 4. Graphical representation of the glycosylated HH<sub>e</sub>- and BB<sub>e</sub>-dimers.**

**Supplementary Fig. 5. Gefitinib alters the cell trafficking of EGFR<sup>L/T/C+KE</sup>.**

**Supplementary Table 1. List of simulated systems with corresponding simulation time and interface strength values.**

**Supplementary Table 2. Primers used in Site-Directed Mutagenesis experiments.**

**Reagents.**

**Supplementary Notes 1-4.**

**References**

**Supplementary Fig. 1. Cartoon and ribbon representations of simulated trimers.**

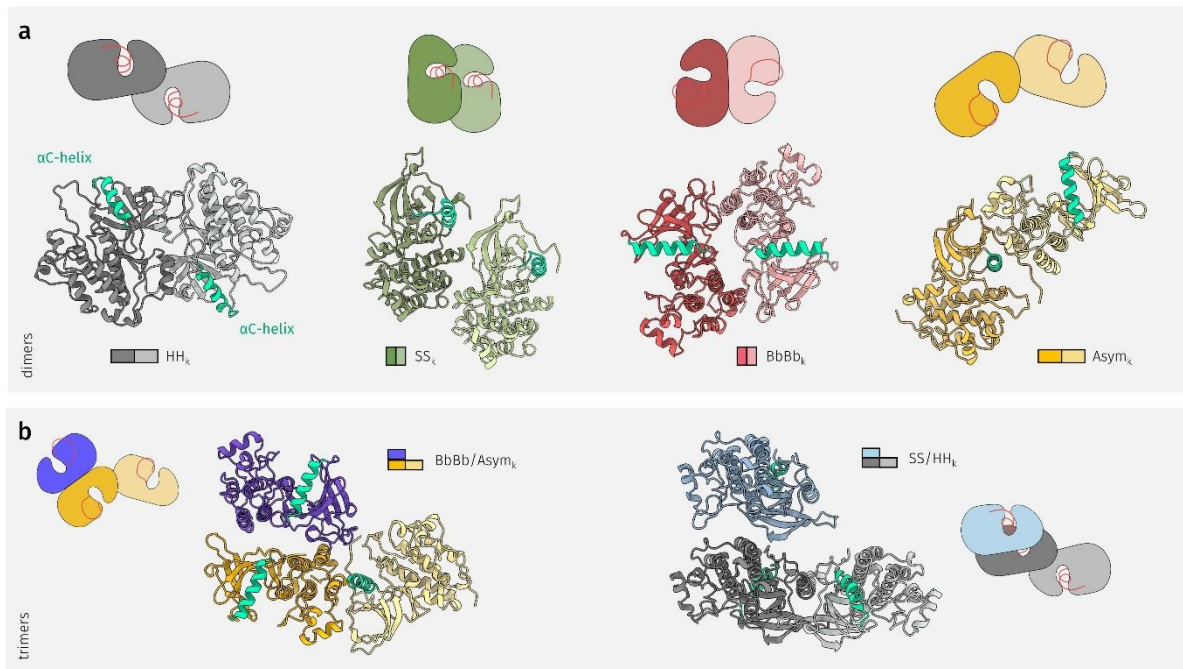

In the cartoon representation of the kinase dimers (**a**, not simulated) and trimers (**b**, simulated), the activation loop is shown in red, distinguishing the active conformation (extended) from the Src-like inactive conformation (with a turn). In the ribbon representation, the αC-helix is highlighted in green to illustrate differences in monomer orientation across the various assemblies. The asymmetric kinase dimer (Asym<sub>k</sub>-dimer) (PDB ID: 2GS6<sup>1</sup>) and head-to-head dimer (HH<sub>k</sub>-dimer) (PDB ID: 5CNO<sup>2</sup>) are present in the main electron density of their respective deposited structures, while the backbone to backbone (BbBb<sub>k</sub>-dimer) (PDB ID: 3VJO<sup>3</sup>) and side-to-side (SS<sub>k</sub>-dimer) (PDB ID: 5CNO) interfaces are observed in crystal lattice contacts of the deposited structures. The SS/HH<sub>k</sub>-trimer (blue/gray) and BbBb/Asym<sub>k</sub>-trimer (purple/orange) are speculative trimeric models constructed to interpret experimental data. The Asym<sub>k</sub>-dimer interface (orange) is mediated by interactions between the αC-helix of the receiver and the C-lobe of the activator. The HH<sub>k</sub>-dimer (gray) is formed primarily through N-to-N lobe contacts, further stabilized by the "electrostatic hook" near the hinge region and the αC/β4 loop. The BbBb<sub>k</sub>-dimer involves mainly N-to-C lobe interactions. The SS<sub>k</sub>-dimer is maintained through contacts between the β2-strand and αD-helix of one monomer and the αE- and αI-helices of the other.

**Supplementary Fig. 2. Key stages in the FLImP data acquisition and analysis process.**

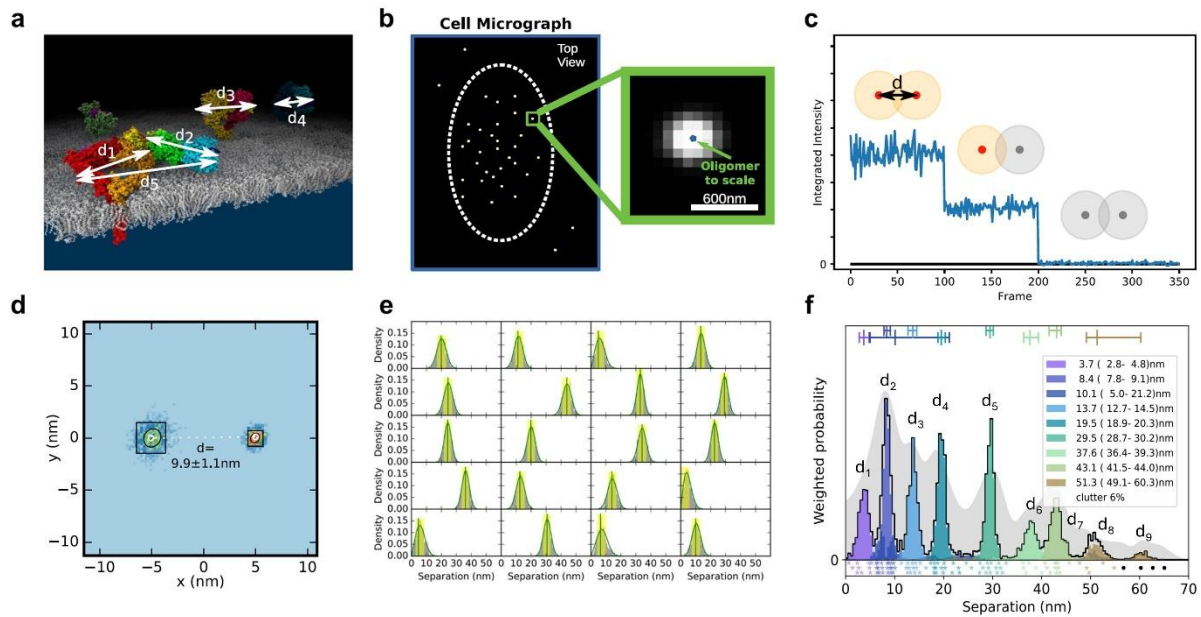

Reproduced from<sup>4</sup>. EGFR is amenable to fluorescent labeling at several sites (**a**), in this case a specific site of the extracellular DIII (the intracellular domain is not depicted); the distances ($d_1$-$d_n$) provide a detailed structural signature. (**b**) When visualized under TIRF microscopy in cells, the size of labeled-EGFR dimers and oligomers is smaller than the diffraction limit of the microscope. Therefore, the fluorescent tags associated to EGFR dimers and oligomers emit light within a diffraction-limited point-spread function (PSF), which is also the microscope image of a single molecule (spots). To resolve the positions of multiple EGFR molecules within a single diffraction limited spot, a video acquisition (FLImP raw data) is taken as the fluorophores photobleach. Single-molecule feature detection and tracking of these videos reveals integrated intensities through time and a subset of these spots which have multistep photobleaching (**c**) is identified (track selection); such single molecules bleach in single steps, and this can be used to estimate the number of fluorophores emitting light (red) in the spot as a function of time. By combining this information with prior knowledge of PSF shape, we can then fit the selected spots through time with a varying sum of Gaussian PSFs model to determine the positions of the emitting fluorophores (FLImP localization fit) with associated uncertainties (**d**). Multiple such measurements in the form of empirical posteriors (**e**) can be pooled (summed) into a FLImP signature of the structure (**f**, gray histogram). The green vertical line in **e** is the mean and the yellow shading represents a 68% confidence interval. Finally, a Bayesian decomposition of the separation measurement set (**e**) is performed to determine the most likely set of unique discrete separations present within the structure (**f**, colored components). In **f** the legend and bars above colored component distributions give the median and most-compact 68% confidence interval for each. Legend also gives median proportion of measurements assigned to clutter<sup>4</sup>.

#### Supplementary Fig. 3. FLImP measurement-dependent oligomer length determination.

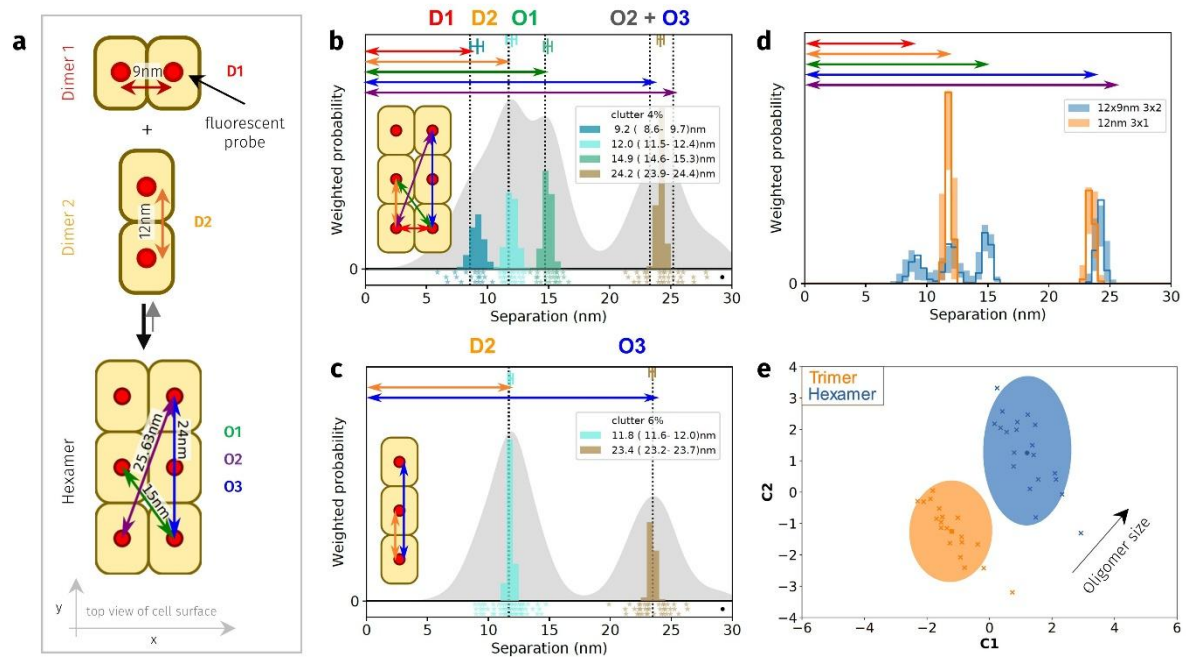

Reproduced from<sup>4</sup>.

(a) Example cartoon of dimer conformers assembling a hexamer. Separations between bound fluorophores (red circles) are: two 1st-order interfaces (D1, D2), short and long diagonals (O1, O2), and 2<sup>nd</sup>-order vertical (O3). (b) FLImP analysis of 100 synthetic pairwise hexamer separations (inset). Sum of posteriors of individual separations between fluorophores (grey background) and abundance weighted probability distributions of individual components of decomposed separation distribution (colored peaks). The area under each peak is weighted according to the estimated proportion of measurements attributed to that peak ("abundance"). Plot legend and bars above colored component distributions give the median and most-compact 68% confidence interval for each. Legend also gives median proportion of measurements assigned to clutter. Stars and dots below show individual separations assigned to peaks (colored stars) or clutter (black dots). (c) As (b) for the homo-trimer formed by the Dimer 2 conformer. (d) Comparisons between decomposed separation probability distributions between datasets. The continuous lines show the marginalized separation posterior for each condition, the sum of the abundance-weighted peaks in (b) and (c). Fluctuations around each continuous line arise from variations derived from FLImP decompositions for 20 bootstrap-resampled datasets to assess errors due to finite number of measurements. e Wasserstein multi-dimensional scale (MDS) analysis of FLImP decompositions. This measures the work needed to convert a decomposed separation set into another, thereby estimating similarities and differences between whole FLImP separation decompositions. Similarities or dissimilarities between the 21 separation sets of different conditions (one main FLImP decomposition plus 20 bootstrap-resampled decompositions) are compared; in this case, for the hexamer, comprised of two conformationally different units (blue) (b) and homo-trimer (orange) (c). The plot axes are components C1 and C2. C1 represents the dimension that captures the largest amount of data variance; C2 represents the second-largest amount of variance orthogonal to C1. The ellipse centers (95% confidence range) mark the positions of the main FLImP decompositions. Crosses mark the positions of individual bootstrap-resampled separation sets.

**Supplementary Fig. 4. Graphical representation of the glycosylated HH<sub>e</sub>- and BB<sub>e</sub>-dimers.**

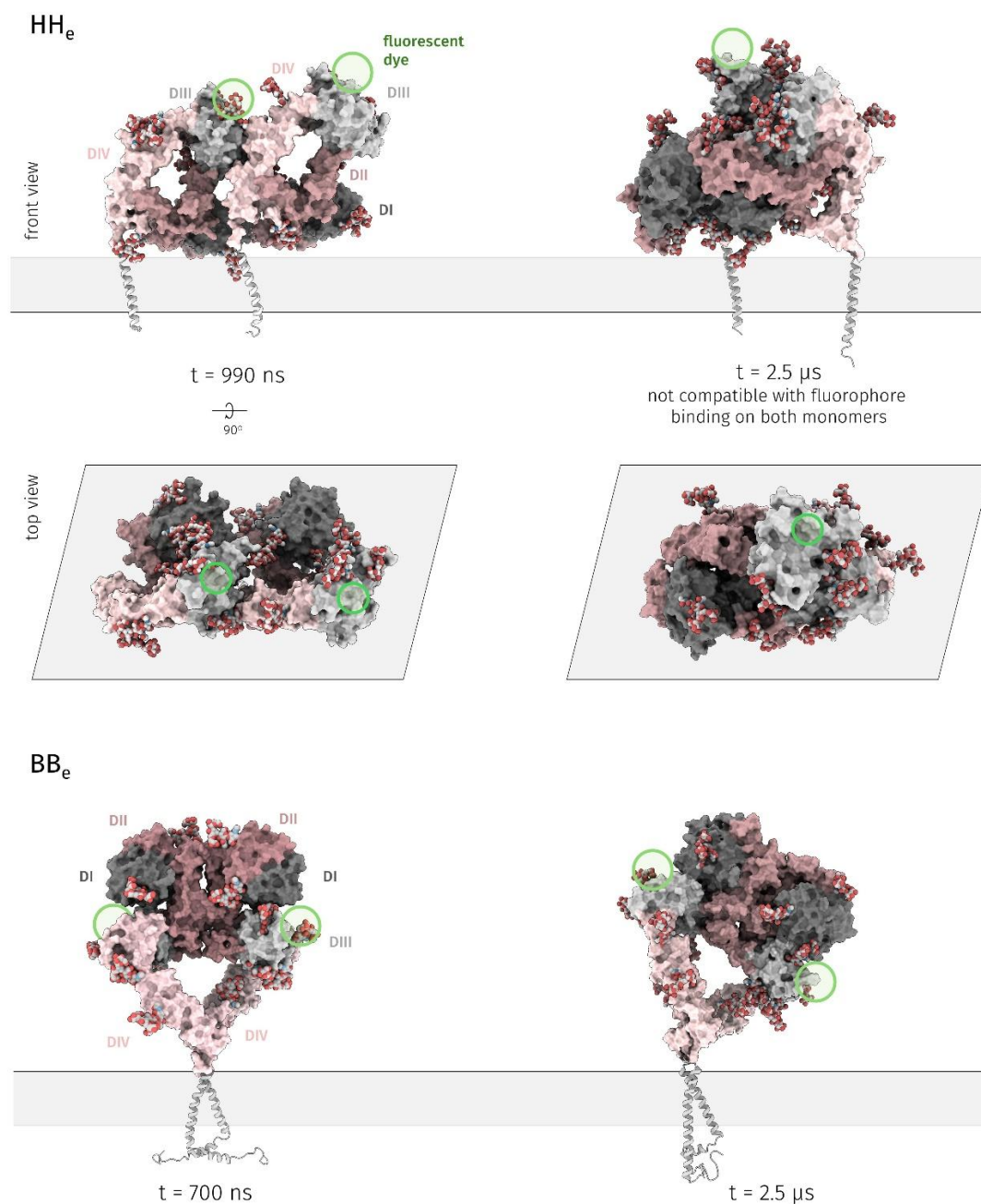

Snapshots from unbiased simulations of the glycosylated, WT ectodomain in the HH<sub>e</sub> (top) and BB<sub>e</sub> (bottom) homodimeric conformations. Glycans are depicted in spheres. The sites where the fluorescent dye is expected to bind are shown in green circles. While the BB<sub>e</sub> stays stable and hence the binding site of the fluorophore dye remains accessible throughout the 2.5 μs simulation, the HH<sub>e</sub> is more flexible, occasionally sampling conformations where the DIII of one monomer “shields” the DIII of the other monomer, which would prevent the fluorophore dye from binding to at least one of the monomers. The simulations did not include the fluorophore dye that is expected to bind to DIII, however, an approximate position is shown in green circles. The simulation details can be found in<sup>4</sup>. Despite the presence of glycans, DIII of one of the monomers is still easily accessible by a fluorophore dye.

**Supplementary Fig. 5. Gefitinib alters the cell trafficking of EGFR<sup>L/T/C+KE</sup>.**

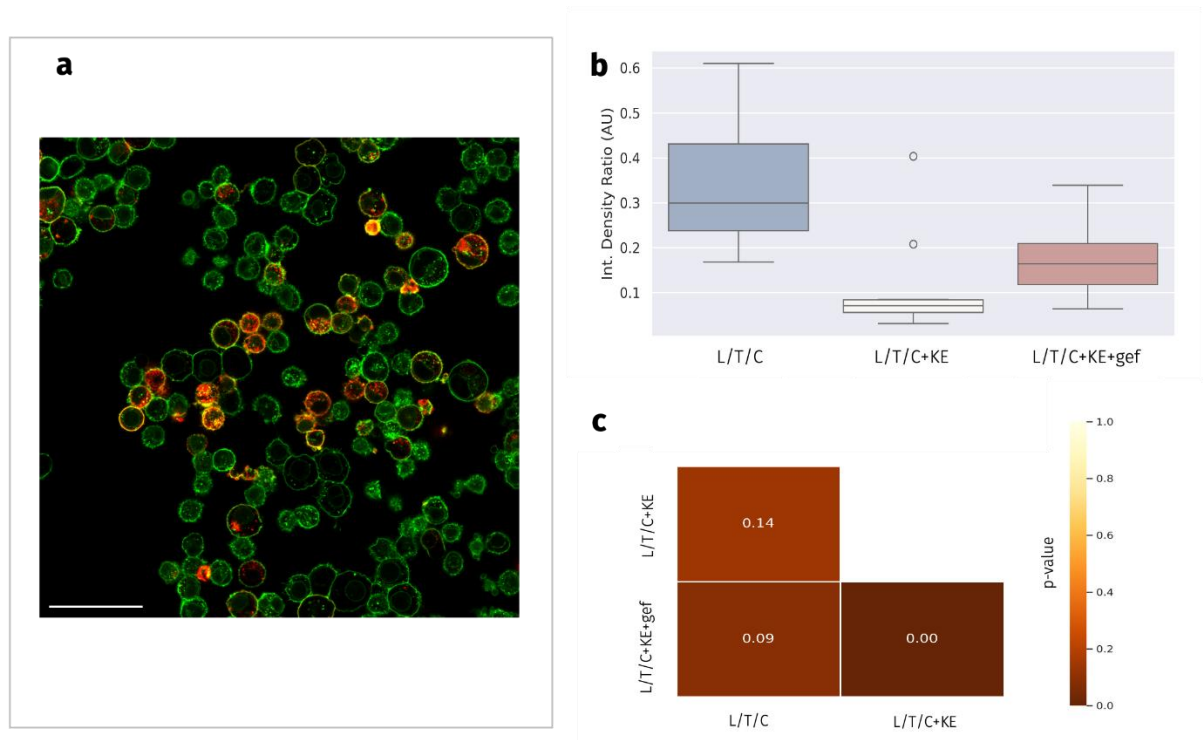

**(a)** Representative image for the analysis in (b) and (c), scale bar = 50 μm. **(b)** Box plots showing the distribution of the ratio between the integrated density of intracellular objects and the integrated density of membrane objects, calculated on a per-cell basis using Fiji ($n \geq 30$). Integrated density is calculated as sum of grayscale pixel values × the number of valid pixels in an object. **(c)** P values for (b), calculated with the `posthoc_ttest` function from the `scikit_posthocs` Python library and the Bonferroni correction for multiple comparisons<sup>5</sup>. These results show that gefitinib has a different effect on the intracellular trafficking of EGFR<sup>L/T/C</sup> and EGFR<sup>L/T/C+KE</sup>. Since EGFR<sup>L/T/C</sup> forms atypical BbBb/Asym<sub>k</sub>-trimers and EGFR<sup>L/T/C+KE</sup> forms canonical Asym<sub>k</sub>-dimers, and because signalling depends on receptor compartmentalization, the differential effect of gefitinib on EGFR<sup>L/T/C</sup> and EGFR<sup>L/T/C+KE</sup> provides preliminary data that the responses of these two variants to the drug might differ and warrants further exploration. Raw data in: https://doi.org/10.5281/zenodo.22129114

**Supplementary Table 1. List of simulated systems with corresponding simulation time and interface strength values.**

| Symmetry | Mutation | Run | Simulation time [μs] | Interface strength |  |
| --- | --- | --- | --- | --- | --- |
| BbBb/Asym<sub>2</sub>-trimer | K970E (KE) | run1 | 2.0 | Asym interface | BbBb interface |
|  |  | run2 | 1.0 | 31.1 | 14.6 |
|  |  | run3 | 2.0 | 27.2 | IC |
|  |  | run4 | 2.0 | 31.3 | IC |
|  | E1005R/D1006K (ED/RK) | run1 | 2.0 | 27.7 | X |
|  |  | run2 | 2.0 | 34.8 | 33.7 |
|  |  | run3 | 2.0 | 38.7 | 24.8 |
|  | T790M | run1 | 2.0 | 34.0 | 12.3 |
|  |  | run2 | 2.0 | 33.8 | 22.9 |
|  |  | run3 | 2.0 | 34.3 | 16.9 |
|  | ΔELREA | run1 | 2.0 | 30.3 | 27.7 |
|  |  | run2 | 2.0 | 25.5 | IC |
|  |  | run3 | 2.0 | 26.9 | IC |
|  | WT | run1 | 2.0 | 19.9 | X |
|  |  | run2 | 1.2 | 38.2 | X |
|  |  | run3 | 2.0 | 37.7 | X |
|  | L858R | run1 | 4.0 | 32.2 | X |
|  |  | run2 | 2.0 | 27.5 | 40.7 |
|  |  | run3 | 2.0 | 26.7 | 26.1 |
|  |  | run4 | 2.0 | 29.6 | 12.9 |
|  |  | run5 | 2.0 | 28.5 | 27.3 |
|  | L858R/T790M (L/T) | run1 | 2.0 | 26.9 | 27.6 |
|  |  | run2 | 2.0 | 40.0 | 27.7 |
|  |  | run3 | 2.0 | 29.0 | 20 |
|  |  | run4 | 2.0 | 26.2 | 26.4 |
|  | L858R/T790M/C797S (L/T/C) | run1 | 2.0 | 28.3 | 23.2 |
|  |  | run2 | 2.0 | 31.7 | 20.5 |
|  |  | run3 | 2.0 | 29.4 | IC |
|  | D770-N771insNPG (InsNPG) | run1 | 2.0 | 28.9 | IC |
|  |  | run2 | 2.0 | 32.1 | X |
|  |  | run3 | 2.0 | 27.7 | 35.8 |
| SS/HH<sub>2</sub>-trimer | K970E (KE) | run1 | 2.0 | HH interface | SS interface |
|  |  | run2 | 4.0 | 45.6 | X |
|  |  | run3 | 4.0 | 41.2 | IC |
|  |  | run4 | 2.0 | 43.1 | 31.3 |
|  | E1005R/D1006K (ED/RK) | run1 | 2.0 | 41.2 | 17.9 |
|  |  | run2 | 2.0 | 45.9 | X |
|  |  | run3 | 2.0 | 50.7 | IC |
|  | T790M | run1 | 2.0 | 44.0 | X |
|  |  | run2 | 2.0 | 39.0 | IC |
|  |  | run3 | 4.0 | 37.4 | IC |
|  |  | run4 | 2.0 | 41.9 | IC |
|  |  | run5 | 2.0 | 46.0 | X |
|  | ΔELREA | run1 | 2.0 | 42.7 | IC |
|  |  | run2 | 2.0 | 39.0 | 24.8 |
|  |  | run3 | 2.0 | 38.1 | IC |
|  | WT | run1 | 2.0 | 46.7 | 22.0 |
|  |  | run2 | 2.0 | 44.3 | 21.5 |
|  |  | run3 | 2.0 | 42.3 | 28.2 |
|  | L858R | run1 | 2.0 | 52.6 | 21.5 |
|  |  | run2 | 4.0 | 41.5 | IC |
|  |  | run3 | 2.0 | 45.0 | IC |
|  | D770-N771insNPG (InsNPG) | run1 | 2.0 | 45.9 | IC |
|  |  | run2 | 2.0 | 33.8 | 12.5 |
|  |  | run3 | 2.0 | 40.7 | 24.4 |
|  | L858R/T790M (L/T) | run1 | 2.0 | 40.4 | 21.5 |
|  |  | run2 | 2.0 | 42.3 | X |
|  |  | run3 | 2.0 | 34.4 | X |
|  | L858R/T790M/C797S (L/T/C) | run1 | 2.0 | 46.0 | 27.2 |
|  |  | run2 | 2.0 | 37.2 | IC |
|  |  | run3 | 2.0 | 45.7 | IC |
|  |  | run3 | 2.0 | 47.6 | IC |

Cell color indicates interface strength, scaled from the darkest red (lowest strength) to the darkest blue (highest strength). Color scales are normalized independently per interface (Asym, BbBb, HH, and SS), as strength values are only directly comparable within the same interface type. X denotes that the interface breaks during the simulation, IC denotes that the interface changes substantially during the simulation without the monomers breaking apart (See Methods section for more details).

### Supplementary Table 2. Primers used in Site-Directed Mutagenesis experiments

Quickchange SDM kit primers:

|  |  |
| --- | --- |
| insNPG EGFR Fw | 5'-GGC TAG CGT GGA CAA CCC CGG CAA TCC TCA CGT GTG CCG CC 3' |
| insNPG EGFR Rv | 5'- GGC GGC ACA CGT GAG GAT TGC CGG GGT TGT CCA CGC TAG CC-3' |
| ΔELREA EGFR Fw | 5'-TCC CGT CGC TAT CAA GAC ATC TCC GAA AGC CA-3' |
| ΔELREA EGFR Rv | 5'-TGG CTT TCG GAG ATG TCT TGA TAG CGA CGG GA-3' |
| L858R EGFR Fw | 5'-GTC AAG ATC ACA GAT TTT GGG CGG GCC AAA CTG CTG GGT GCG-3' |
| L858R EGFR Rv | 5'-CGC ACC CAG CAG TTT GGC CCG CCC AAAATC TGT GAT CTT GAC-3' |
| T790M EGFR Fw | 5'-CCT CCA CCG TGC AGC TCA TCA TGC AGC TCA TGC CCT TCG GC-3' |
| T790M EGFR Rv | 5'-GCC GAA GGG CAT GAG CTG CAT GAT GAG CTG CAC GGT GGA GG-3' |
| C797S EGFR Fw | 5'-CAT GCC CTT CGG CAG CCT CCT GGA CTA-3' |
| C797S EGFR Rv | 5'-TAG TCC AGG AGG CTG CCG AAG GGC ATG-3' |
| K970E EGFR Fw | 5'-GGG GGT CTC GGG CCA TCT CGG AGAATT CGA TGA TC-3' |
| K970E EGFR Rv | 5'-GAT CAT CGA ATT CTC CGA GAT GGC CCG AGA CCC CC-3' |
| E1005R/D1006K EGFR Fw | 5'-CAC CAC GTC GTC CAT CTT TCT TTC ATC CAT CAG GGC ACG GTA GAA GTT G-3' |
| E1005R/D1006K EGFR Rv | 5'-CAA CTT CTA CCG TGC CCT GAT GGA TGA AAG AAA GAT GGA CGA CGT GGT G-3' |

NEB SDM kit primers:

|  |  |
| --- | --- |
| E1005R/D1006K EGFR Fw | 5'-GAT GGA TGA ACG CAA AAT GGA CGA CGT G-3' |
| E1005R/D1006K EGFR Rv | 5'-AGG GCA CGG TAG AAG TTG G-3' |

### Reagents

Antibodies and reagents were purchased as follows: Anti-EGFR Affibody® Molecule (Abcam ab95116; RRID: AB\_11156238), Anti-EGFR EgB4 nanobody (gift from Paul M.P. van Bergen en Henegouwen at Division of Cell Biology, Science Faculty, Department of Biology, Utrecht University, Utrecht 3584 CH, The Netherlands), Anti-EGFR (D38B1, Cell Signaling Technology (4267; RRID: AB\_2246311), Anti-beta-Actin Monoclonal Antibody, HRP Conjugated (13E5 Cell Signaling Technology 5125; RRID: AB\_1903890), Anti-Phospho EGF Receptor (Tyr1068, D7A5 Cell Signaling Technology 3777; RRID: AB\_2096270), Anti-Phospho EGF Receptor (Tyr1173, 53A5, Cell Signaling Technology 4407; RRID: AB\_331795), Anti-EGFR (R & D Systems, AF231; RRID: AB\_355220), Anti-EGFR (1005, Santa Cruz Biotechnology), Anti-Phospho-Akt (Ser473, D9E, Cell Signaling Technology 4060; RRID: AB\_2315049), Anti-Akt (Cell Signaling Technology 9272; RRID: AB\_329827), Anti-Phospho p44/42 (ERK1/2, Thr202/Tyr204, D13.14.4E, Cell Signaling Technology 4370; RRID: AB\_2315112), Anti-p44/42 (ERK 1/2 Cell Signaling Technology 9102; RRID: AB\_330744), Anti-Phospho-STAT5 (Tyr694, C11C5, Cell Signaling Technology 9359; RRID: AB\_823649), Anti-STAT5 (D2O6Y, Cell Signaling Technology 94205; RRID: AB\_2737403), Anti-

Phospho AMPKa (Thr 172, 40H9, Cell Signaling Technology 2535; RRID: AB\_331250), Anti- AMPKa (D5A2, Cell Signaling Technology 5831; RRID: AB\_10622186), Anti-Phospho p38 (Thr 180/Tyr182, clone 36, BD Transduction Laboratories™ 612288; RRID: AB\_399605), Anti-p38 (Cell Signaling Technology 9218; RRID: AB\_10694846), Anti-VEGF Receptor 2 (D5B1, Cell Signaling Technology 9698; RRID: AB\_11178792), Anti-mouse IgG HRP (Jackson ImmunoResearch 715-035-150; RRID: AB\_2340770), Anti-rabbit IgG-HRP (Jackson ImmunoResearch 711-035-152; RRID: AB\_10015282), Gefitinib (Sigma-Aldrich, SML1657) or Iressa (Tocris Bioscience, 3000), Recombinant Murine EGF (Peprotech, 315-09), Recombinant Murine IL-3 (Peprotech, 213-13), 4% Paraformaldehyde (EMGrade, Electron Microscopy Sciences, 157-4), 25% Glutaraldehyde (Grade I, Sigma G5882), FluoSpheres™ (carboxylate modified, 0.1 µm, infrared (715/755), Invitrogen F8799).

#### **Supplementary Note 1 — Modelling of the kinase trimers**

Supplementary Fig. 1 shows a graphical representation of the kinase trimers we have simulated as well as the dimers that are discussed throughout the text, even if they were not simulated. The Asym<sub>k</sub>-dimer and HH<sub>k</sub>-dimer have been resolved and reported in the main electron density of their respective deposited structures, while the BbBb<sub>k</sub>-dimer and SS<sub>k</sub>-dimer are observed in the crystal lattice of deposited structures. In contrast, the HH/SS<sub>k</sub>-trimer and BbBb/Asym<sub>k</sub>-trimer are speculative trimeric models we have constructed to interpret experimental data. Further validation details for these trimeric models can be found in our previous work<sup>4</sup>.

A caveat of our *in silico* modeling is that our protocol simulates preformed trimers in different arrangements, thereby neglecting any mutation-driven effects on the structure and dynamics of the underlying monomers—effects that could potentially prevent dimer and/or trimer formation altogether. However, the FLImP separation data provide evidence for trimer formation, confirming that trimers do assemble under these conditions. This lends some support to our modeling assumption of preformed trimers.

**BbBb/Asym<sub>k</sub>-trimer.** The simulations, which included an Asym<sub>k</sub>-dimer, namely the BbBb/Asym<sub>k</sub>-trimer, were based on the dimers seen in the crystal structure of EGFR bound to an ATP analogue (PDB entry 2GS6<sup>1</sup>). This type of dimer is mediated through interactions of the αC-helix of the receiver with the C-lobe of the activator, and its formation has been associated with the promotion of transphosphorylation<sup>1</sup>. After removing the bound ligand, we introduced the L858R and T790M single-point mutations, the ΔELREA deletion, the InsNPG insertion, the double L858R/T790M mutation, the L858R/T790M/C797S triple mutation, as well as the non-naturally occurring K970E mutation, to the WT structure using MODELLER<sup>6</sup>. Each monomer of the dimer consisted of the S695-G983 sequence (in the numbering with the 24-aa tag). For this reason, we did not simulate the E1005R/D1006K double mutant, as modelling part of the highly flexible exon 25 C-terminal tail to the Asym<sub>k</sub>-dimer, for which there is no data on how it should fold, would introduce a high degree of uncertainty to the model.

The simulations, which included a BbBb<sub>k</sub>, namely the BbBb/Asym<sub>k</sub>-trimer, were based on the dimers seen in the crystal packing of PDB entry 3VJO<sup>3</sup>. The BbBb<sub>k</sub>-dimer described in<sup>1</sup>, is formed mainly through N-to-C lobe interactions. We removed the co-crystallised ligand from both monomers, and we built missing atoms using MODELLER. Each monomer of the dimer consisted of the G696-G1022 sequence. As in the case of the Asym<sub>k</sub>-dimer, we introduced the ΔELREA, L858R, T790M, K970E, InsNPG, L858R/T790M, and L858R/T790M/C797S mutations with

MODELLER. In the case of the InsNPG, even though there is an available crystal structure for this mutant, we still modelled the mutant following the same MODELLER protocol for consistency, but we then compared the structure of each monomer of the resulting modelled dimer to the crystal structure of InsNPG in complex with a covalent inhibitor (PDB entry 4LRM<sup>7</sup>) to ensure that the overall conformation of each monomer and the environment around the point of the mutation are in accordance with the experimentally derived structure.

Through a series of site-specific mutations and MD simulations, we showed in our previous work that monomers can dock into an Asym<sub>k</sub>-dimer to form a BbBb/Asym<sub>k</sub>-trimer<sup>4</sup>. To construct the BbBb/Asym<sub>k</sub>-trimer that we used in the simulations, we superimposed one of the BbBb<sub>k</sub> monomers onto one of the Asym<sub>k</sub> monomers. Given the sequence of the Asym<sub>k</sub> monomers, the E1005R/D1006K mutation was introduced only in the BbBb<sub>k</sub> monomer of the BbBb/Asym<sub>k</sub>-trimer. For the rest of the systems, the mutation was introduced to all three monomers.

The HH<sub>k</sub>, described in<sup>8</sup>, is formed mainly through N-to-N lobe interactions and further stabilized by the “electrostatic hook” located near the hinge region of the kinase domain and near the αC/β4 loop. The simulations, which included the HH<sub>k</sub> were based on the primary dimer resolved in PDB entry 5CNO<sup>2</sup>. Each monomer of the dimer consisted of the N700-E1015 sequence.

**SS/HH<sub>k</sub>-trimer.** The co-occurrence of the HH<sub>k</sub>-dimer and SS<sub>k</sub>-dimer in the crystal lattice of the V948R EGFR (PDB entry 5CNO) made us speculate about the existence of a “zig-zag” tetramer in cellular contexts, comprising a HH<sub>k</sub> subunit and two monomers attached to it in a SS<sub>k</sub> arrangement. In this proposed tetramer model, the two monomers of the HH<sub>k</sub> are found in an Src-like inactive conformation, while the two monomers attached to the HH<sub>k</sub> can adopt an active or inactive conformation, as their αC-helix and A-loop are not part of the SSI interaction interface. This flexibility suggests that monomers can attach to a central HH<sub>k</sub>-dimer and form a SS/HH<sub>k</sub>-trimer or a SS/HH<sub>k</sub>-tetramer. The SS/HH<sub>k</sub>-trimer that we used in the simulations was constructed by superimposing one of the SS<sub>k</sub> monomers onto one of the HH<sub>k</sub> monomers according to the crystal structures.

### Supplementary Note 2 — MD simulation protocol

For consistency, we applied the same simulation protocol to all the different dimers and trimers. In particular, for the unbiased MD simulations we performed, each simulated system was parameterized using the CHARMM36m force field<sup>9</sup>. The protonation states of the residues at pH 7.4 were determined using the Proteinprepare functionality of PlayMolecule suite<sup>10</sup>, which maintained the usual charge states for all residues. We then solvated each system using modified TIP3P CHARMM water molecules<sup>11</sup> in a dodecahedral box, while Na<sup>+</sup> and Cl<sup>-</sup> ions were added to neutralize the system's charge and reach a final concentration of 0.15 M. The GROMACS v.2022.5 engine<sup>12</sup> was used for all the simulations. Prior to the production simulations, the energy of each system was minimized using steepest-descent energy minimization. After minimization, the initial velocities for the atoms were then taken from a Maxwell distribution at 300 K. We then simulated the system for 5 ns in the NVT ensemble using a velocity rescaling thermostat<sup>13</sup> and harmonic position restraints on protein heavy atoms (1000 kJ mol<sup>-1</sup> nm<sup>-2</sup>), followed by 10 ns in the NPT ensemble under constant pressure (1 bar) using the Berendsen barostat<sup>14</sup>, followed by further 5 ns using the Parrinello-Rahman barostat<sup>15</sup>. All bond lengths to hydrogen atoms were constrained using the LINCS algorithm<sup>16</sup>, while van der Waals interactions were treated with a cut-off distance of 10 Å. Electrostatic interactions were computed using the particle mesh Ewald

method<sup>17</sup> with a direct sum cut-off distance of 10 Å and a Fourier spacing of 1.6 Å. After equilibration, we ran single or multiple independent replicas of unbiased simulations for each system. A summary of the different simulations that we ran, and their corresponding length, can be found in Supplementary Table 1. In cases where any of the monomers were breaking apart or showing major interface changes, we interrupted the simulation before the end of it, hence the difference in simulation time for some systems. The list of all the simulated systems and the simulation time of each can be found in Supplementary Table 2.

#### **Supplementary Note 3 — Interface strength analysis**

To analyze the strength of the different interfaces, we used the Python library *mdciao*<sup>18</sup> (Supplementary Table 2). The calculation of the interface strength is based on the time-traces of residue-residue distances, from which we derive the contact frequencies and distance distributions. The average strength of the interface corresponds to the sum of all the average numbers of neighbours. In other words, a higher average strength suggests that the interface gains contacts upon mutation, or loses contacts with respect to the WT if the interface strength of the mutant is lower than that of the WT. For the definition of the interfaces, we used a 4 Å cut-off to consider all residues at the interface of each dimer within this cut-off for analysis. Given that the residue pairs at the interface whose distances and frequencies we monitor over the MD trajectory are computed only at the beginning of the simulation, in cases where the interface changes substantially and the initial residue pairs no longer describe the same interface, we do not report the value of the strength, but we label the event as an interface change (IC).

The simulations that we ran to quantify the strength of the different interfaces produce three qualitatively distinct outcomes: a numeric strength value, a fracture event (X), or an IC. These outcomes are informative, not interchangeable, and therefore, we present the outcomes of each simulation explicitly.

An X event is a real observation about the low end of the strength distribution where the interface broke. When a system produces a mix of numeric values and X events, averaging only the numeric values yields an upward-biased estimate of strength, because the broken runs are silently excluded. The X events are telling us that the true distribution has a lower tail, which we are not seeing. X events are therefore reported explicitly.

IC events are different in nature. An IC event does not mean the interface was weak, but rather that the interface is transformed so fundamentally that the strength metric loses its physical meaning. It is a statement that the question itself became invalid for that simulation, not that the answer was low. IC events are therefore excluded from the strength distribution and are reported explicitly from both the numeric values and the X events. For systems where IC is the only observed outcome, no strength distribution is shown at all because none can be honestly claimed.

#### **Supplementary Note 4 — Tumor growth analysis**

Tumor volumes were modeled with a linear mixed-effects model using log1p-transformed tumor volume, which preserves zero-volume observations that a plain logarithm would exclude. Fixed effects included time (day), treatment system (system), and their interaction (day × system), and the model included mouse-specific random intercepts together with random slopes for time to account for within-mouse correlation across repeated measurements. Models were fit by maximum likelihood. The primary test compared the full model (day × system) against a reduced model omitting the system terms (day only) using a likelihood-ratio test, to assess whether growth trajectories differed overall across systems. Pairwise contrasts between systems were then evaluated at two prespecified timepoints, day 48 and day 124.

Using all available longitudinal measurements (n = 759 observations from 25 mice across 5 systems; day range 0–124), the mixed-effects model showed an overall difference in tumor trajectories across systems (LRT: $\chi^2(8) = 48.81$, $p = 6.92 \times 10^{-8}$, \*\*\*). Pairwise contrasts at day 48 and day 124 are reported as model-estimated log-differences and fold-change ratios, with associated p-values and significance levels, in Extended data Table 1.
