## Extended data: Figures and tables for "Disrupting aberrant EGFR catalytic trimers reverses T790M gefitinib resistance"

### CONTENT:

**Extended data Fig. 1. Photobleaching correlation spectroscopy of receptor oligomerization.**

**Extended data Fig. 2. FLImP data are consistent with a shared oligomer architecture.**

**Extended data Fig. 3. Effect of HH<sub>e</sub>-dimer conformational fluctuations on ectodomain separations stabilized by kinase interactions.**

**Extended data Fig. 4. FLImP decompositions in the 0-70 nm range.**

**Extended data Fig. 5. (a): Probe binding to the BB<sub>e</sub>-HH<sub>k</sub>-dimer in ΔELREA and T790M oligomers is sterically hindered. (b): The KE mutation alters oligomer assembly kinetics and diffusion. (c) Gefitinib induces a conformational change in EGFR<sup>T790M+KE</sup> oligomers.**

**Extended data Fig. 6. Western blots and quantification.**

**Extended data Fig. 7. *Ex vivo* evaluation of the excised tumours based on GFP fluorescence and calliper measurement to compare tumor growth in different cohorts.**

**Extended data Fig. 8. Immunohistochemistry of xenograft tumor sections.**

**Extended data Fig. 9. Top: Significance and enrichment scores for the microarray data.**

**Extended data Table 1: (a) Kaplan Meir pairwise comparison of time to progression for T790M, gefitinib, and the KE mutation. (b) Pairwise comparisons of tumor growth between treatment systems at days 48 and 124.**

### References

### Extended data Fig. 1. Photobleaching correlation spectroscopy of receptor oligomerization.

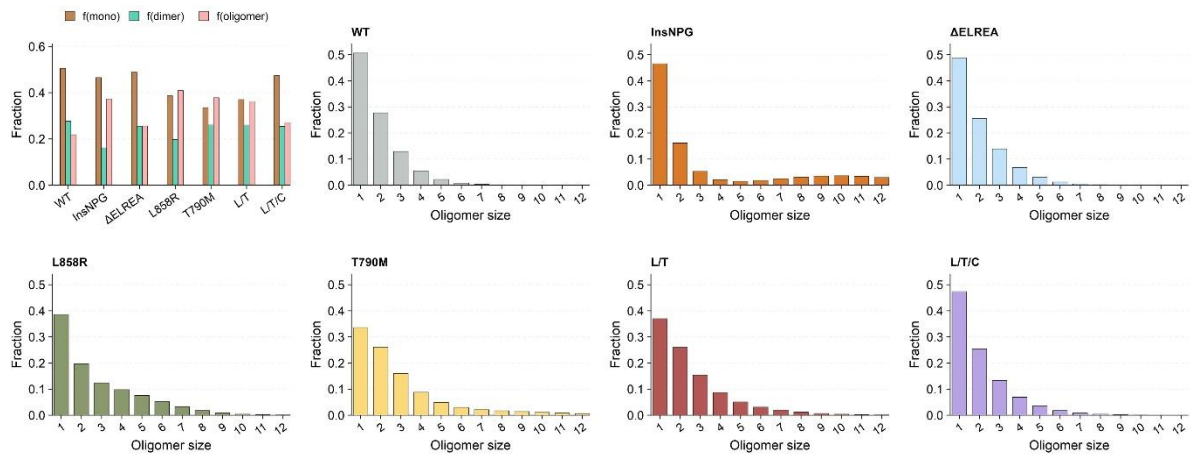

Molecular-normalized fraction of receptors in oligomer species measured in CHO cells expressing ~100,000 receptor copies per cell. Cells were treated with 100 nM Alexa 488-Affibody, and the oligomer fraction was determined by photobleaching imaging correlation spectroscopy, as previously shown<sup>1</sup>. The average fraction of dimers and oligomers corresponding to these data is also shown. The lower and upper limits in the fraction of oligomers are estimates depending on the size of oligomers. We considered sizes from tetramers up to oligomers with 20 protomers.

**Extended data Fig. 2. FLIMP data are consistent with a shared oligomer architecture.**

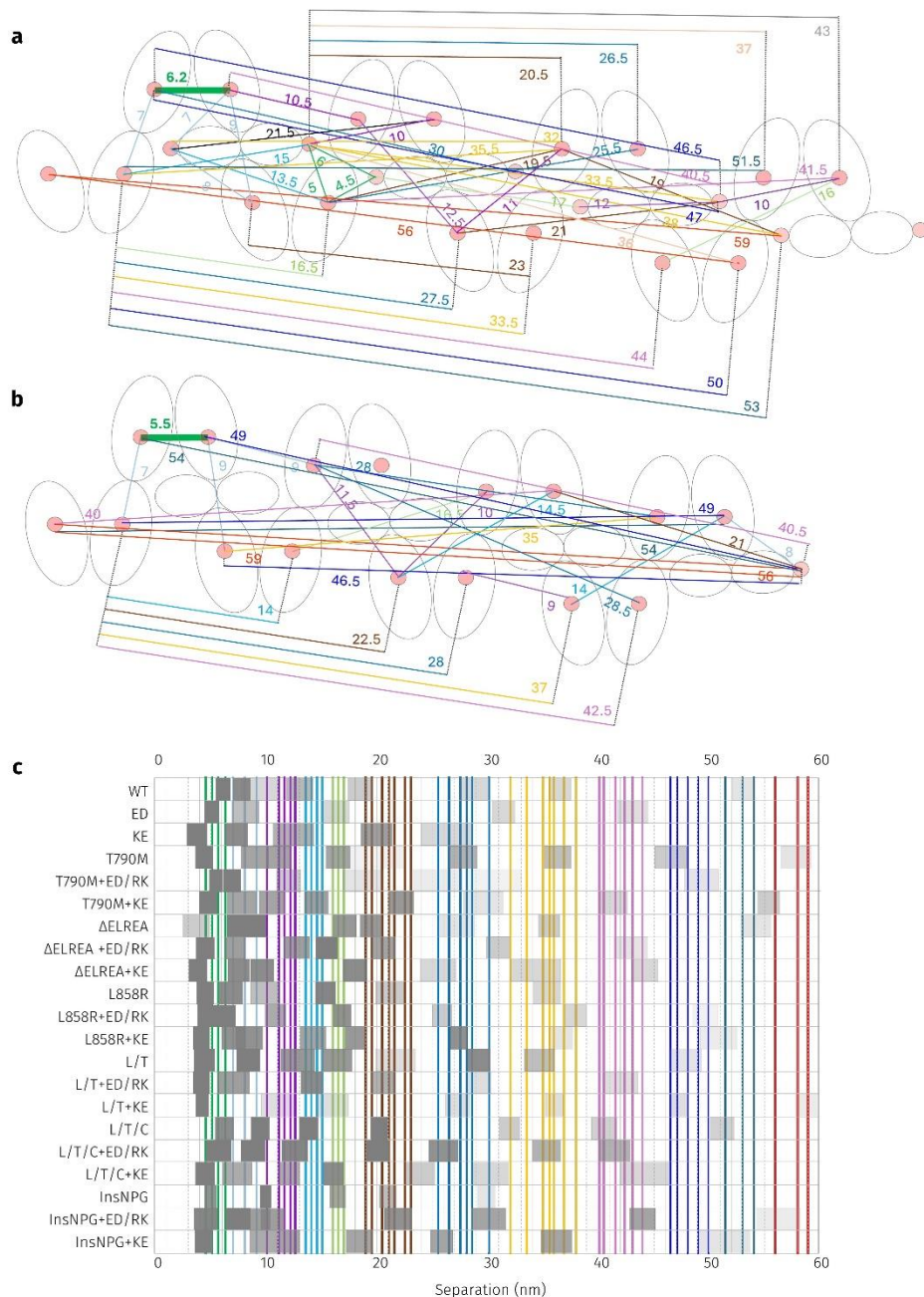

Top view of the EGFR oligomer<sup>2</sup>. Separations between the two DIII labels in the HH<sub>e</sub>-dimers are 6.2 nm in the extended model (a) and 5.5 nm in the compacted model (b). These values represent the two maxima in the section of the distribution of possible separations in HH<sub>e</sub>-dimers that are compatible with oligomer assembly (see Extended data Fig. 3 and Supplementary Fig. 4). Separations between DIII labels (pink circles) in both models are indicated by continuous lines. Lines and numbers are color-coded to aid identification. **(c)** Table showing the FLIMP separations determined for EGFR<sup>WT</sup> and its variants in the extended 0-60 nm range, including native forms and those with the KE or ED/RK unnatural mutations, as described in the main text (decomposed data in Extended Fig. 4). The width of the gray rectangles represents 1SD in these separations, and their intensity reflects the relative height of the peak components within each condition. Vertical colored lines superimposed on the table of separations denote the separations measured in the models in (a) and (b). All separations predicted were aligned closely with the experimental data, supporting the conclusion that the relevant variants share a common oligomer architecture.

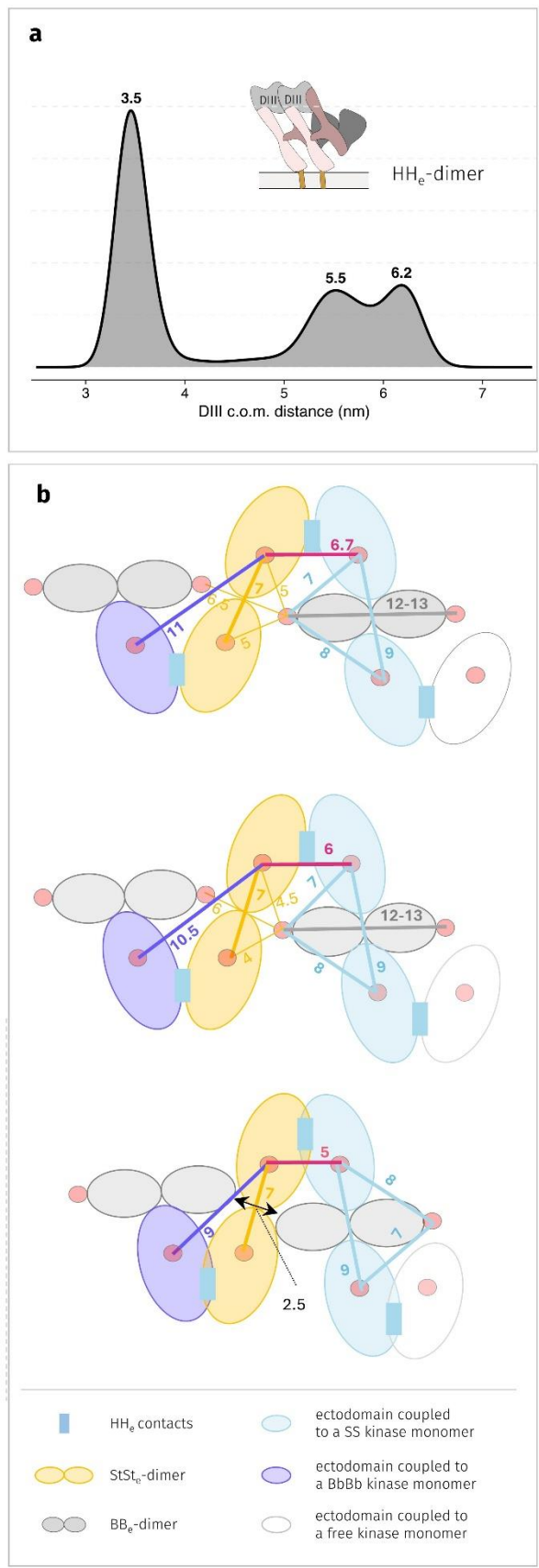

**Extended data Fig. 3. Effect of HH<sub>e</sub>-dimer conformational fluctuations on ectodomain separations stabilized by kinase interactions. (a)** Center of mass (c.o.m.) distance between domains III (DIIIs) of an HH<sub>e</sub>-dimer. The bimodal distribution represents the c.o.m. distance over a 2.5 μs long MD simulation of glycosylated dimers embedded in a membrane via their transmembrane (TM) domains. Details can be found in<sup>2</sup>. The conformational fluctuations of the HH<sub>e</sub>-dimer are significant when compared with the $\sim 2$ nm resolution of FLImP. **(b)** Model of the EGFR oligomer (top view). Separations between the two DIIIs labels in the HH<sub>e</sub>-dimers are set to 6.7 nm (top), 6.0 nm (middle), and 5.0 nm (bottom). The distribution mode below 4 nm produces a steric clash between ectodomains (black arrow, bottom oligomer). Separations between 5-6.7 nm therefore define the allowable range in HH<sub>e</sub>-dimers compatible with their participation in oligomer assembly. We estimated the theoretical uncertainties in separations between ectodomains introduced by conformational HH<sub>e</sub>-dimer fluctuations by measuring distances between ectodomain DIII c.o.m. across three oligomers. Separations underpinned by the SS/HH<sub>k</sub>-trimer (blue) are independent of DIII separations in the HH<sub>e</sub>-dimers. The $\sim 7$ nm separation of the StSt<sub>e</sub>-dimer (orange) arises from the value measured in the model. This $\sim 7$ nm separation is also independent of DIII separations in the HH<sub>e</sub>-dimers. The StSt<sub>e</sub>-dimer was previously experimentally validated<sup>2</sup>; because a crystal structure is unavailable to anchor a simulation, the only non-experimental uncertainty for this separation came from spatial measurement errors within the model; done to the scale 1 cm = 1 nm this error is small ($\sim 0.1$ nm). For display purposes, we used 1 nm as nominal error in main text Fig. 3a.

**Extended data Fig. 4. FLImP decompositions in the 0-70 nm range.**

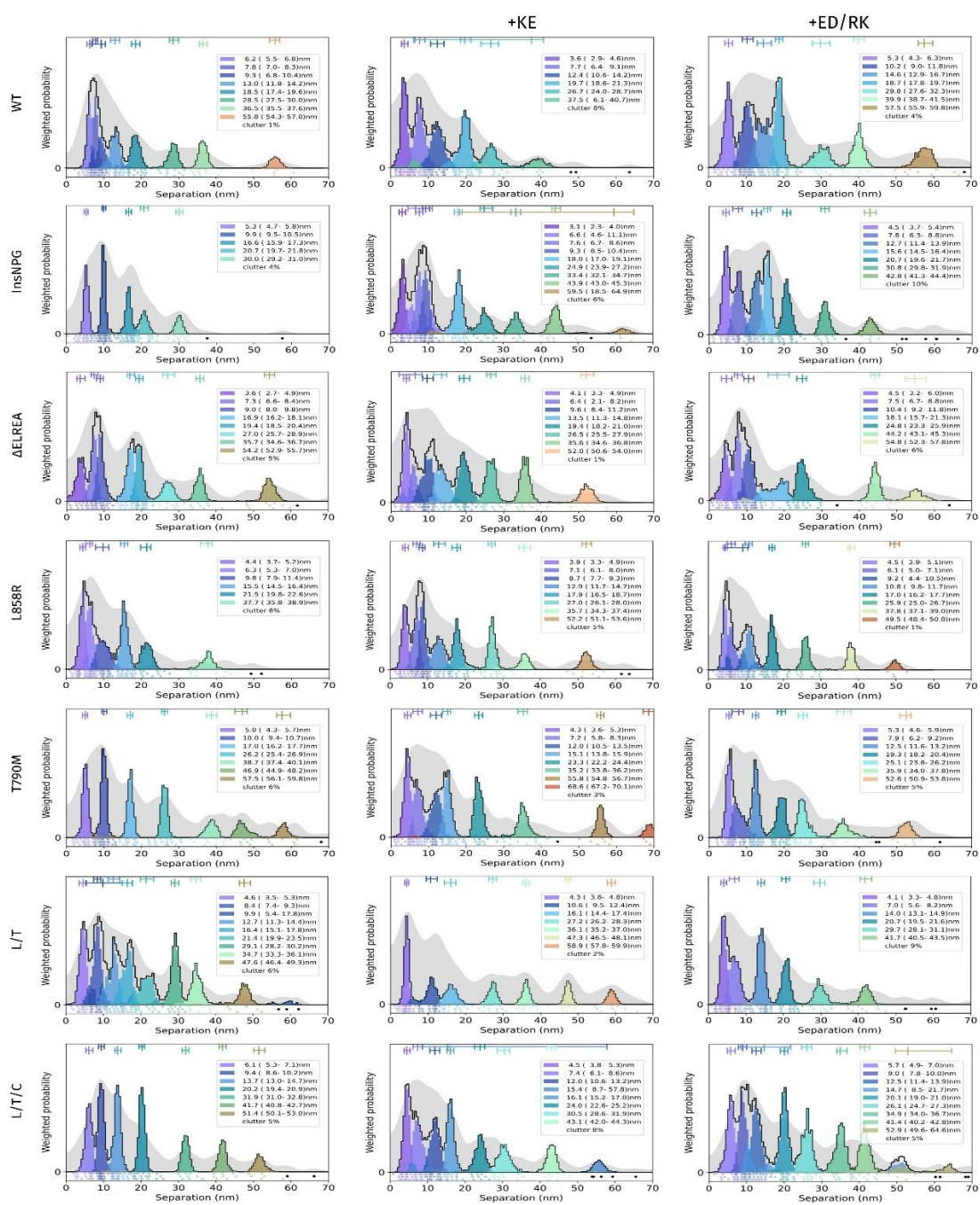

FLImP analysis of 100 separation probability distributions with the smallest uncertainty between Affibody-CF640R pairs in the range 0-70 nm: Sum of posteriors of individual separations between fluorophores (gray background) and abundance-weighted probability distributions of individual components of decomposed separation distribution (colored peaks). The continuous black line depicts the marginalized separation posterior for each condition, which is the sum of the abundance-weighted peaks. The plot legend and bars above colored component distributions give the median and most-compact 68% confidence interval for each. The legend also gives the median proportion of measurements assigned to clutter. Stars and dots below show individual separations assigned to peaks (colored) or clutter (black). The full-width uncertainty of each of the 100 empirical posterior separations is noted in nanometers. The typical number of empirical separations per condition was ~1,000 - 2,000.

**Extended data Fig. 5. (a): Probe binding to the BB<sub>e</sub>-HH<sub>k</sub>-dimer in ΔELREA and T790M oligomers is sterically hindered. (b): The KE mutation alters oligomer assembly kinetics and diffusion. (c) Gefitinib induces a conformational change in EGFR<sup>T790M+KE</sup> oligomers.**

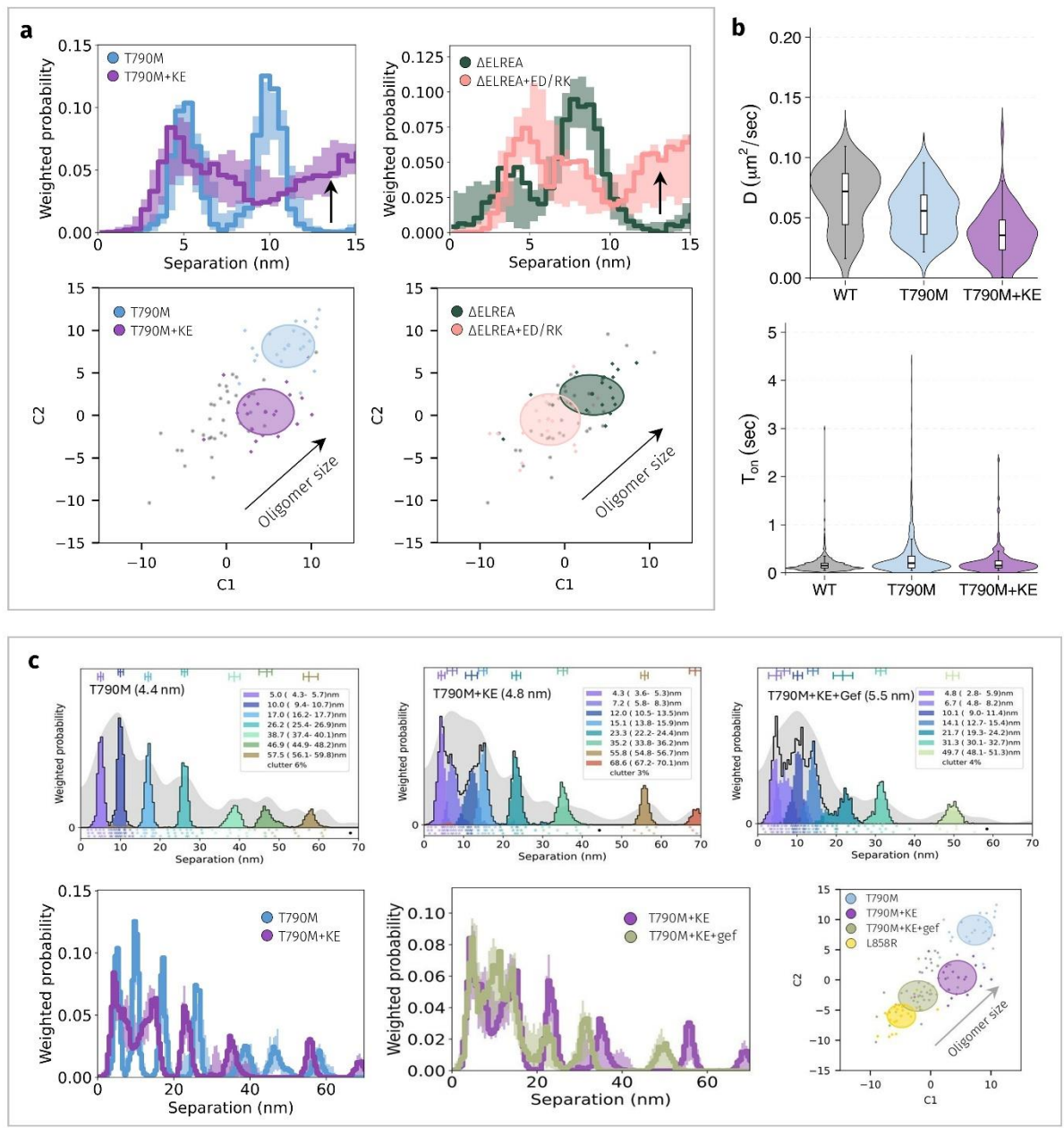

**(a) Probe binding to the BB<sub>e</sub>-HH<sub>k</sub>-dimer in ΔELREA and T790M oligomers is sterically hindered.** Comparisons of decomposed separation probability distributions between datasets. Continuous lines show the marginalized separation posterior, the sum of the abundance-weighted peaks. Fluctuations around each continuous line arise from variations derived from FLImP decompositions for 20 bootstrap-resampled datasets to assess errors due to finite number of measurements. The data show an increase in separations consistent with a doubly labelled BB<sub>e</sub>-HH<sub>k</sub>-dimer (12-13 nm) when the KE and ED/RK mutations are applied to EGFR<sup>T790M</sup> and EGFR<sup>ΔELREA</sup>, respectively. Bottom: Wasserstein multidimensional scaling (MDS) analysis of the above FLImP decompositions. The MDS metric maps similarities and differences between 21 separation sets of different conditions (one main FLImP decomposition plus 20 bootstrap-resampled decompositions). The plot axes are components C1 and C2. C1 represents the dimension that captures the largest amount of variance in the data, while C2 represents the second-largest amount of variance that is orthogonal to C1. Uncertainty ellipses represent $1\sigma$. For more details, see Supplementary Fig. 7. As shown, the recovery of the fluorescence signal from the BB<sub>e</sub>-HH<sub>k</sub>-dimer is not accompanied by an increase in oligomer size. The latter rules out the additional recruitment of BB<sub>e</sub>-HH<sub>k</sub>-dimers and shows that the absence of the 12-13 nm peak was due to steric hindrance. **(b): The KE mutation alters oligomer assembly kinetics and diffusion.** Two-color single particle tracking comparing diffusion rates (D) and T<sub>on</sub> values as previously described<sup>3</sup>. The latter measures the duration of the interaction between tracking particles. The data show significant differences in D and T<sub>on</sub> value distributions with and without adding the KE mutation (p=0.0003 and p=0.01, respectively). Data acquired over at least 30 cells, over 3 independent biological replicates. **(c) Gefitinib induces a conformational change in EGFR<sup>T790M+KE</sup> oligomers.** FLImP decompositions (top) and comparisons between the FLImP separation decompositions (bottom left, middle) (details of how these comparisons are made in (a)). In contrast to the comparison between EGFR<sup>T790M+KE</sup> and EGFR<sup>T790M</sup>, the differences between EGFR<sup>T790M+KE</sup> and EGFR<sup>T790M+KE</sup> + gef in the ~0-30 nm range are borderline significant, falling within the errors reported by bootstrapping with resampling. However, a Wasserstein multi-dimensional scale (MDS) analysis of FLImP decompositions to compare oligomer size (described in (a)) shows that gefitinib significantly decreases the size of EGFR<sup>T790M+KE</sup> oligomers (bottom right). This reduction brings them to a size comparable to that of EGFR<sup>L858R</sup>, which assembles the shortest oligomers in this study (Main text, Fig. 3f). The short oligomers assembled by EGFR<sup>T790M+KE</sup> + gefitinib reflect a lack of engagement of the BB<sub>e</sub>-HH<sub>k</sub>-dimers, which are the components responsible for oligomer growth<sup>2</sup>. This is unsurprising because gefitinib preferentially binds the active conformation of the kinase domain, which is incompatible with the autoinhibited BB<sub>e</sub>-HH<sub>k</sub>-dimers.

### Extended data Fig. 6. Western blots and quantification.

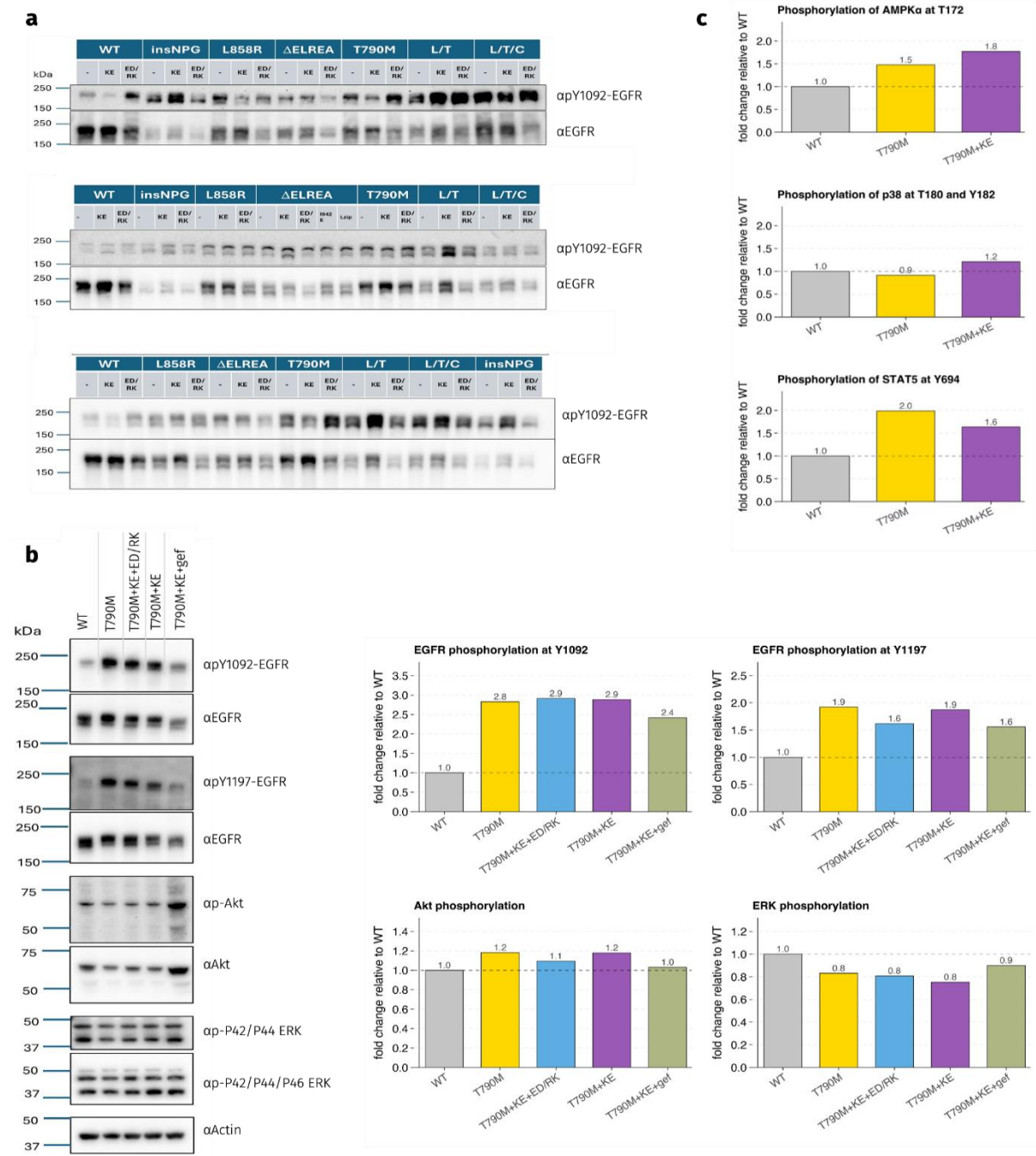

**(a)** Western blot results ($n = 3$) showing ligand-independent phosphorylation at EGFR tyrosine residue 1092 of oncogenic variants transiently transfected in CHO cells (see methods). Blots were probed with Anti-Phospho-EGF Receptor (Tyr1092) and an equal combination of Anti-EGF Receptor antibodies plus Anti rabbit IgG-HRP. Quantification of these blots is shown in Fig. 3g in the main text **(b)** (Left) Western blot analysis of the phosphorylation EGFR variants at tyrosine residue 1092 and tyrosine residue 1197; the variants were stably expressed in Ba/F3 cells in the absence of IL3 and with or without $1\mu\text{M}$ gefitinib treatment. Western blots also show phosphorylation of endogenous Akt and p44/42 MAPK (ERK 1/2). (Right) Quantification of these blots, relative to cells expressing WT EGFR, is shown below. **(c)** Quantification of Western blots in Fig 5c showing phosphorylation of endogenous AMPKα, p38, and STAT5 in the absence of both growth factor stimulation and IL3 in stably transfected Ba/F3 cells. Phosphorylation levels of these proteins in cells expressing EGFR variants are shown relative to cells expressing WT EGFR.

**Extended data Fig. 7. *Ex vivo* evaluation of the excised tumours based on GFP fluorescence and calliper measurement to compare tumor growth in different cohorts.**

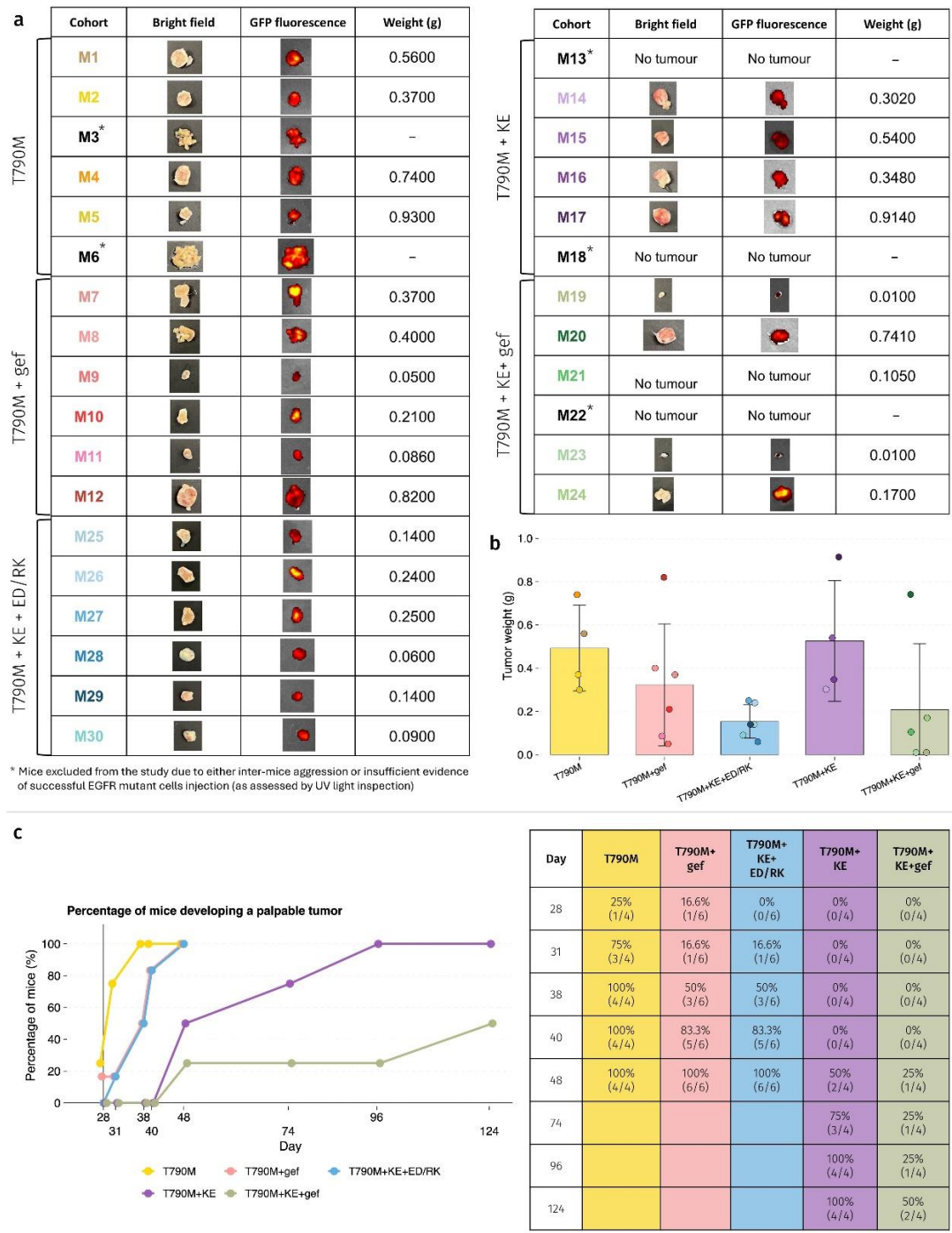

Male NOD scid gamma (NSG) mice were used to establish tumors in their flanks (see Methods) and tumor growth was monitored using callipers. N=6 animals per cohort. IVIS imaging of all animals at the experimental endpoint showed retained GFP fluorescence, as an indication of tumor growth. Harvested tumors from those animals with established tumors were weighed (*right panel*), thereafter imaged under daylight (*left panel*) and blue light illumination to reflect GFP fluorescence (*middle panel*). **(b)** Comparison of tumor weights among different cohorts; error bars are 1SD. **(c)** Cumulative plot of the percentage of mice that had developed a palpable tumor over the course of the study.

**Extended data Fig. 8. Immunohistochemistry of xenograft tumor sections, related to main text Figs. 4 and 5.**

**a EGFR**

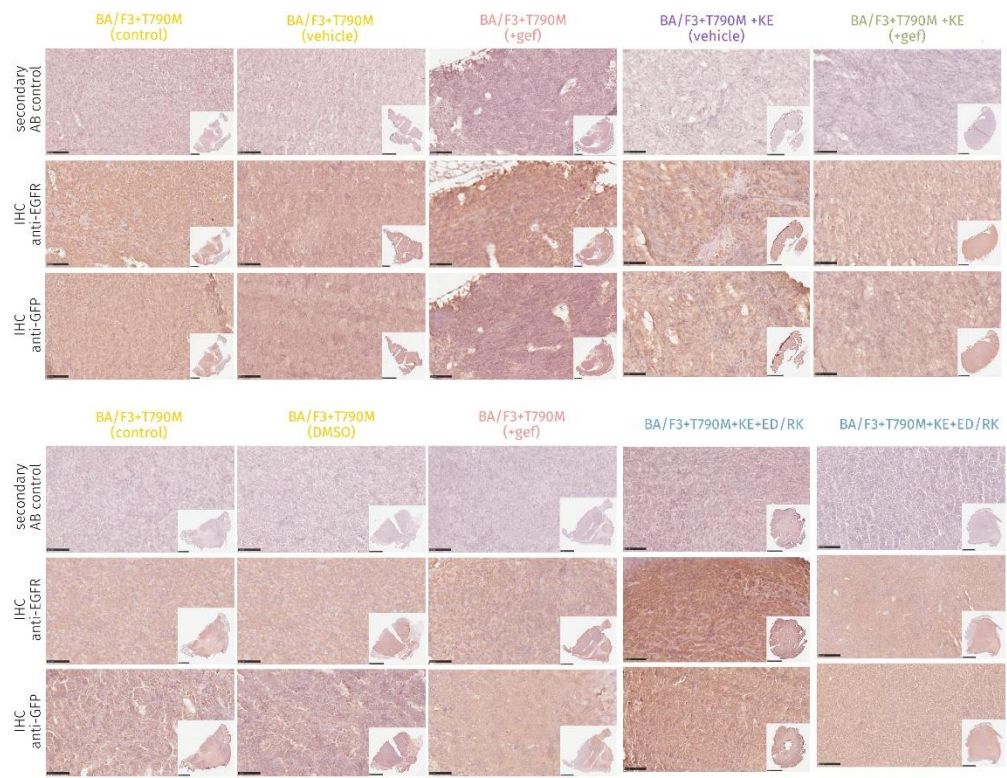

**b VEGFR2**

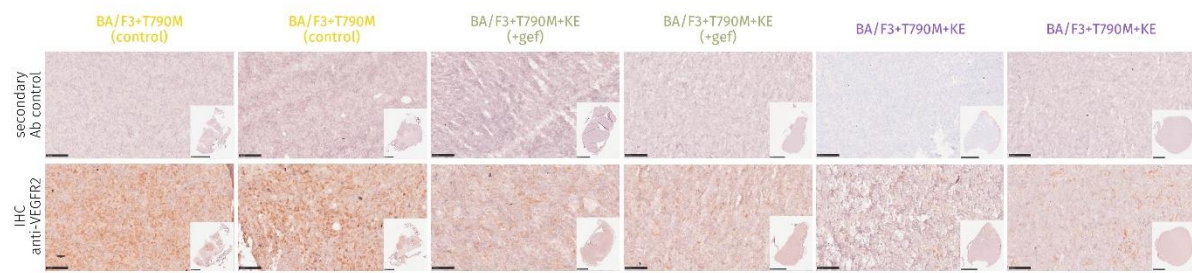

**(a)** Immunohistochemistry staining of tumor sections to support *in vivo* tumor growth using an anti-GFP antibody. These stains show retained EGFR expression at the endpoint of the study using a pan anti-EGFR antibody. **(b)** Immunohistochemistry staining of tumor sections to evaluate the tumor vasculature as an indication of angiogenesis using an anti-VEGFR antibody. All scale bars represent 100 $\mu\text{m}$.

**Extended data Fig. 9. Top: Significance and enrichment scores for the microarray data.**

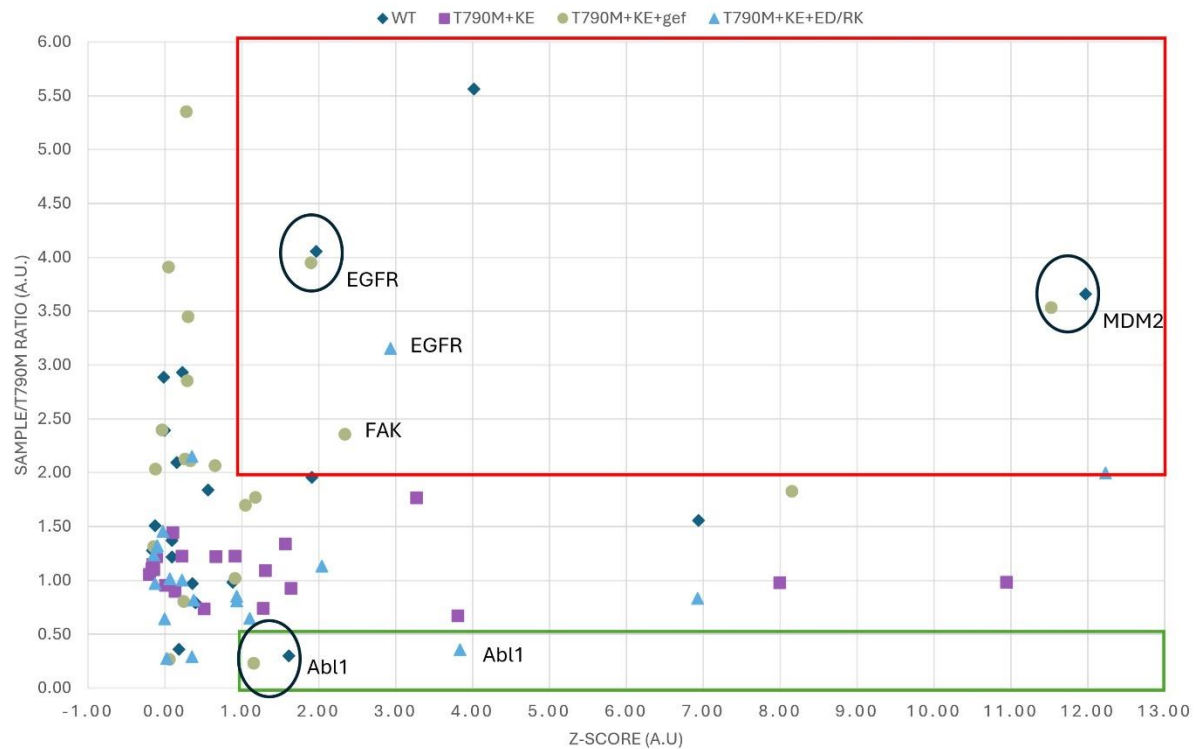

The Z score for the raw value of $x_i$ is defined as: $z_i = (x_i - \langle x \rangle / \sigma_x)$, where $\langle x \rangle$ is the mean of all the sample values, and $\sigma_x$ is the standard deviation of these values, as described in<sup>4</sup>. Fold change was calculated per gene against expression in the T790M dataset. Hits were highlighted if Z-score >1 (expression > 1σ above assay average) and fold change > 2 (red box) or <0.5 (green box). Pairs of hits where scores closely align between WT and T790M+KE+gef datasets are highlighted by a dark blue circle.

#### Extended data Table 1:

##### (a) Pairwise comparisons of tumor growth between treatment systems at days 48 and 124.

| Day | System 1 | System 2 | log diff | fold change | se | z | p value | Significance |
| --- | --- | --- | --- | --- | --- | --- | --- | --- |
| 48 | T790M | T790M+KE+gef | 5.9560 | 386.0621 | 0.4997 | 11.9186 | 9.468E-33 | *** |
| 48 | T790M | T790M+KE | 5.0952 | 163.2355 | 0.5214 | 9.7729 | 1.471E-22 | *** |
| 48 | T790M+KE+ED/RK | T790M+KE+gef | 3.9924 | 54.1838 | 0.4406 | 9.0609 | 1.293E-19 | *** |
| 48 | T790M+gef | T790M+KE+gef | 3.8052 | 44.9354 | 0.4406 | 8.6362 | 5.812E-18 | *** |
| 48 | T790M+KE | T790M+KE+ED/RK | -3.1316 | 0.0436 | 0.4650 | -6.7345 | 1.645E-11 | *** |
| 48 | T790M+gef | T790M+KE | 2.9444 | 18.9997 | 0.4650 | 6.3320 | 2.420E-10 | *** |
| 48 | T790M | T790M+gef | 2.1508 | 8.5915 | 0.5272 | 4.0798 | 4.507E-05 | *** |
| 48 | T790M | T790M+KE+ED/RK | 1.9636 | 7.1250 | 0.5272 | 3.7248 | 1.955E-04 | *** |
| 48 | T790M+KE | T790M+KE+gef | 0.8608 | 2.3651 | 0.4336 | 1.9851 | 4.714E-02 | * |
| 48 | T790M+gef | T790M+KE+ED/RK | -0.1872 | 0.8293 | 0.4715 | -0.3969 | 6.914E-01 | ns |
| 124 | T790M+KE | T790M+KE+gef | 8.9722 | 7880.9501 | 0.5983 | 14.9952 | 7.890E-51 | *** |

Log differences are fitted contrasts on the log<sub>10</sub>p scale; fold change is exp(log difference). A positive log difference means a larger predicted tumor volume in system 1 than in system 2. The day-124 contrast is limited to T790M+KE versus T790M+KE+Gef, the only systems still measured at that day. T790M, T790M+Gef, and T790M+KE+ED/RK end at day 48, and the quadratic spline is not interpreted beyond their last observation. Significance codes: ns, $p \geq 0.05$; \*, $p < 0.05$; \*\*, $p < 0.01$; \*\*\*, $p < 0.001$. More details in Supplementary Note 4

##### (b) Kaplan Meir pairwise comparison of time to progression for T790M, gefitinib, and the KE mutation.

| System A | System B | Mice in A | Progressions in A | Mice in B | Progressions in B | Standard p-value | Significance |
| --- | --- | --- | --- | --- | --- | --- | --- |
| T790M | T790M+KE+gef | 4 | 4 | 5 | 2 | 0.0029 | ** |
| T790M | T790M+KE | 4 | 4 | 4 | 4 | 0.0069 | ** |
| T790M | T790M+gef | 4 | 4 | 5 | 5 | 0.0400 | * |
| T790M+KE | T790M+KE+gef | 4 | 4 | 5 | 2 | 0.0483 | * |

Time to progression was compared across four systems: T790M, T790M+gef, T790M+KE, and T790M+KE+gef. The three T790M+KE+gef mice that had not progressed by days 119–124 were kept as censored observations. The comparison therefore uses 18 mice: 15 that progressed and 3 that were still progression-free at the end of follow-up. Each pair was tested with a two-sided log-rank test, the standard comparison for Kaplan–Meier curves. The standard p-value is the usual $\chi^2$ result.

#### Extended Data References

1 Ciccotosto, G. D., Kozer, N., Chow, T. T., Chon, J. W. & Clayton, A. H. Aggregation distributions on cells determined by photobleaching image correlation spectroscopy. *Biophys J* **104**, 1056-1064, doi:10.1016/j.bpj.2013.01.009 (2013).
2 Iyer, R. S. *et al.* Drug-resistant EGFR mutations promote lung cancer by stabilizing interfaces in ligand-free kinase-active EGFR oligomers. *Nat Commun* **15**, 2130, doi:10.1038/s41467-024-46284-x (2024).
3 Zanetti-Domingues, L. C. *et al.* The architecture of EGFR's basal complexes reveals autoinhibition mechanisms in dimers and oligomers. *Nat Commun* **9**, 4325, doi:10.1038/s41467-018-06632-0 (2018).

4 Brideau, C., Gunter, B., Pikounis, B. & Liaw, A. Improved statistical methods for hit selection in high-throughput screening. *J Biomol Screen* **8**, 634-647, doi:10.1177/1087057103258285 (2003).
